# Neuronal distribution across the cerebral cortex of the marmoset monkey (*Callithrix jacchus*)

**DOI:** 10.1101/385971

**Authors:** Nafiseh Atapour, Piotr Majka, Ianina H. Wolkowicz, Daria Malamanova, Katrina H. Worthy, Marcello G.P. Rosa

## Abstract

Using stereological analysis of NeuN-stained sections, we investigated neuronal density and number of neurons per column throughout the marmoset cortex. Estimates of mean neuronal density encompassed a greater than threefold range, from >150,000 neurons/ mm^3^ in the primary visual cortex to ~50,000 neurons/ mm^3^ in the piriform complex. There was a trend for density to decrease from posterior to anterior cortex, but also local gradients, which resulted in a complex pattern; for example, in frontal, auditory and somatosensory cortex neuronal density tended to increase towards anterior areas. Anterior cingulate, motor, premotor, insular and ventral temporal areas were characterized by relatively low neuronal densities. Analysis across the depth of the cortex revealed greater laminar variation of neuronal density in occipital, parietal and inferior temporal areas, in comparison with other regions. Moreover, differences between areas were more pronounced in the supragranular layers than in infragranular layers. Calculations of the number of neurons per unit column revealed a pattern that was distinct from that of neuronal density, including local peaks in the posterior parietal, superior temporal, precuneate, frontopolar and temporopolar regions. These results suggest that neuronal distribution in adult cortex result from a complex interaction of developmental/ evolutionary determinants and functional requirements.

## Introduction

There has been renewed recent interest in obtaining precise estimates of neuronal density in the primate cerebral cortex, sparked by attempts to establish organizational principles that link cytoarchitecture, development and function (e.g. Cahalane et al. 2012; Charvet et al. 2015; Hilgetag et al. 2016). Nonetheless, for many areas, it remains surprisingly difficult to obtain comprehensive information about this basic characteristic of cortical organization.

To date, the only comprehensive data sets that cover most of the primate cortex have been obtained using the isotropic fractionator method (Collins et al. 2010, 2011; Turner et al. 2016). However, this method only allows a very approximate correlation between cell density values and areal boundaries, and results in irretrievable loss of potentially important information about the neuronal density in different layers. Moreover, there is evidence that this technique carries several potential sources of imprecision, which become apparent when the results are compared with those obtained using stereological methods (reviewed by Charvet et al. 2015). On the other hand, studies which included a combination of stereological methods and expert determination of cytoarchitectural areas have not yet fully sampled the primate cortex, with the number of areas studied ranging from a few (Rockel et al. 1980; Hustler et al. 2005; Sherwood et al. 2007; Turner et al. 2016) to a few dozen (Dombrowski et al. 2001; Hilgetag et al. 2016). In addition, most of these studies have relied on the Nissl stain, which tends to slightly overestimate the number of neurons (Hilgetag et al. 2016). In summary, no study to date has obtained a complete picture of how neuronal density varies between the areas of the cerebral cortex of any primate species.

Here we report on neuronal density in the cortex of the marmoset monkey (*Callithrix jacchus*), a New World monkey which is attracting increasing interest as a model species for studies of neural systems that are particularly differentiated in the primate brain (Solomon and Rosa 2014; Mitchell and Leopold 2015; Majka et al. 2016; Miller et al. 2016; Eliades and Miller 2017; Hagan et al. 2017; Oikonomidis et al. 2017; Okano and Kishi 2017; Silva 2017). The neuronal density was assessed from the surface to the bottom of layer 6 in each of the 116 cytoarchitectural areas currently recognized in the cortex of this species (Paxinos et al. 2012), through stereological analysis of sections stained for a neuron-specific marker (NeuN; Mullen et al. 1992). This resulted in estimates which are both finer-grained, and more accurately mapped to the functional boundaries of areas, than previously available. Based on this data set, and computational measurements of cortical thickness, we were also able to evaluate variations in the numbers of neurons per equal area column, as well as the total numbers of neurons in the different areas. The entire set of digital images used in this study is shared through a freely accessible web site (http://www.marmosetbrain.org/cell_density), which will hopefully enable future re-analyses in light of the rapidly evolving knowledge about the marmoset brain.

## Materials and Methods

Tissue was obtained from the right hemispheres of one female (CJ167) and two male (CJ173, CJ800) marmosets (*Callithrix jacchus*). The animals were sexually mature young adults (26, 24, and 32 months of age, respectively). Each animal was also used for neuroanatomical tracing experiments unrelated to this study. The entire set of sections through the cortex, and the results of the anatomical tracing experiments, can be visualized at the Marmoset Brain Architecture Project web site (http://www.marmosetbrain.org/injection). Given the uniform quality of the stain throughout the brain, case CJ167 provided the most extensive data set, which allowed us to obtain detailed estimates for every cortical area. Data from cases CJ173 and CJ800 were obtained in a reduced set of areas, with the objective of estimating the degree of inter-individual variation. All procedures were approved by the Monash University Animal Ethics Experimentation Committee, which also monitored the health and wellbeing of the animals throughout the experiments. The animals had no veterinary record of serious or chronic health conditions.

### Histology

The methods used for tissue preparation have been described in detail previously (Atapour et al. 2017). Briefly, at the conclusion of the anatomical tracing experiment, the marmosets were overdosed with sodium pentobarbitone sodium (100 mg/kg) and transcardially perfused with 0.1 M heparinized phosphate buffer (PB; pH 7.2), followed by fixative (4 % paraformaldehyde in 0.1 M phosphate buffer). The brains were dissected and kept in the same fixative solution for 24 h, followed by immersion in fixative solutions containing increasing concentrations of sucrose (10%, 20% and 30%).

Frozen 40 μm coronal sections were obtained using a cryostat. Every fifth section was set aside for NeuN staining. Other series were stained for cytochrome oxidase activity (Silverman and Tootell 1987) and myelin (Gallyas 1979) to help delineate laminar and areal boundaries. An additional series was left unstained, for the analysis of fluorescent tracers, and another was kept in reserve. For NeuN staining, free-floating sections were incubated in blocking solution (10% normal horse serum and 0.3% Triton-X100 in PBS) for 1 h at room temperature, and then incubated with the primary antibody for NeuN (1:800, MAB377, clone A60, Merck Millipore) at 4°C for 42-46 h. Secondary antibody (1:200, PK-6102, Vectastain elite ABC HRP kit, Vector Laboratories) was applied for 30 min at room temperature, followed by avidin/biotin interaction and DAB (DAB Peroxidase Substrate Kit SK-4100, Vector Laboratories) staining. Following mounting, the sections were examined under a Zeiss AxioImager light microscope at high magnification, under an oil immersion objective, to assess uniform penetration of the antibodies. In all 3 animals there were no gaps or regions of paler staining in the sections used for analysis, including at a focal plane corresponding to the mid-thickness of the sections. The NeuN-stained sections were then scanned using Aperio Scanscope AT Turbo (Leica Biosystems) at 20× magnification, with the focus set to the mid-thickness plane. This provided a resolution of 0.50 μm/pixel, while ensuring that the vast majority of neurons showing nuclear staining were visible in the images used for analysis.

Within each coronal section, we delineated each of the cortical areas proposed by Paxinos et al. (2012), using the cytoarchitectural criteria illustrated in that atlas, as well as myelo-and chemoarchitectural criteria detailed in previous publications from our group (e.g. Palmer and Rosa 2006a, b; Burman and Rosa 2009; Reser et al. 2009, 2013; Rosa et al. 2009). We then undertook computerized reconstruction of the entire hemisphere (Majka et al. 2016), which converted the series of sections into a 3-dimensional volume. The process of assigning of each voxel within this volume to a cortical area was constrained by the expert delineation of the cytoachitectural areas.

### Estimating neuronal density

Four sections spanning the rostro-caudal extent of each architectonic area in approximately equal steps, were selected for analysis. Within each of these sections we selected a columnar-shaped counting strip, based on location (as far from the estimated boundaries with other areas as possible), the absence of staining artifacts (such as detachments, folds or injection sites), and the absence of large blood vessels (which could introduce significant gaps in the estimates). Whenever possible, we also tried to select regions of low curvature, but this was not always possible due to the locations of some of the areas; thus, in some areas the cell density profiles across the cortical depth show variations (Supplementary Fig. S1), as expected from the differential expansion and compression of layers according to the convexity of the cortex (Hilgetag and Barbas 2006). Moreover, for areas located in the frontal and occipital pole (cytoarchitectural area 10, and the primary visual cortex, V1) we avoided regions where the sections were tangent or near-tangent to the surface. Finally, in topographically organized visual areas the analysis included samples from the foveal, intermediate and peripheral representations (Rosa et al. 1997; Rosa and Elston 1998; Rosa and Tweedale 2000, 2005; Chaplin et al. 2013), whereas in somatosensory and motor areas they included samples from different body part representations (Krubitzer and Kaas 1990; Burman et al. 2008). The original high-resolution images of the counting strips used in the analyses are freely available for download at http://www.marmosetbrain.org/cell_density.

Using standard features of the Aperio Image Scope software, square counting frames (150×150 μm) were aligned across the cortical thickness, with the top of the uppermost box aligned with the interface between layers 1 and 2 (Fig. 1). Due to the step change in neuronal density which characterizes the boundary between layers 1 and 2, and the fact that layer 1 was typically thinner than the height of the frames used for most of the quantification, estimates of neuronal density in layer 1 were obtained separately, using smaller counting frames which only included parts of this layer. These frames were rectangular, having the same width of the frames used for quantifying layers 2-6 (i.e. 150 μm parallel to the layers), but variable heights (75-125 μm) depending on the area. In addition, given that the use of the NeuN stain revealed that white matter neurons (Mortazavi et al. 2016, 2017) were ubiquitous, the placement of counting frames was continued below layer 6, to account for such neurons. The number of counting frames varied depending on the thickness of the cortex in each section. Thus, to allow alignment of the values obtained in different locations of the same area according to the same proportional depth, we used a procedure that considered depth 0 as the top of layer 2, and depth 1.0 the bottom of layer 6, estimated visually in the sections. Values of neuronal density assigned to depths >1.0 corresponded to sampling frames centered on the white matter; as explained in Results, these were not used in the calculations of mean neuronal density.

**Figure 1.**
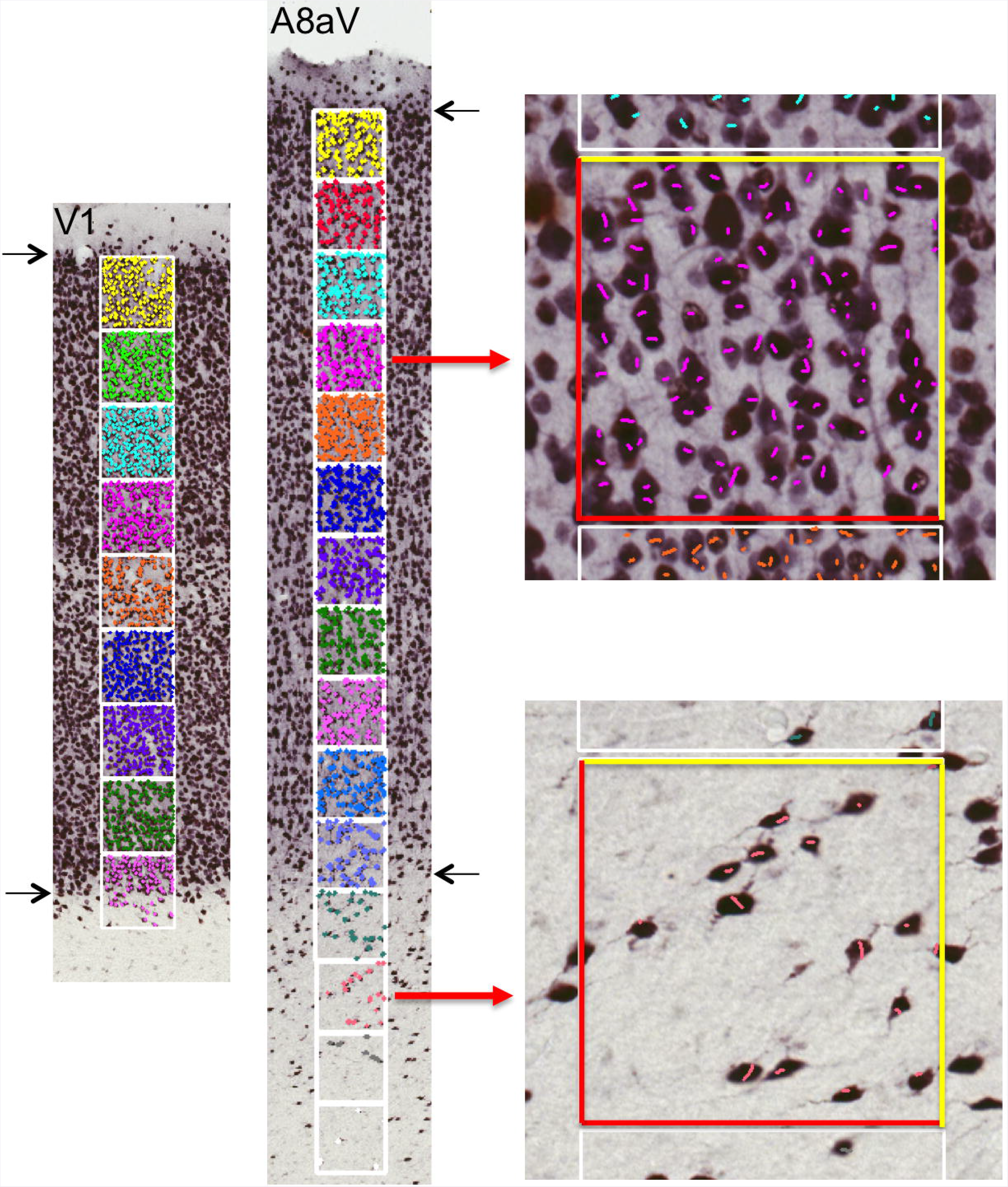
Sampling method for stereological analysis. The placement of square counting frames (0.15×0.15 mm) is illustrated on example NeuN-stained sections from primary visual cortex (V1) and ventral subdivision of area 8a (A8aV), covering the depth of the cortex from layer 2 to the white matter subjacent to layer 6. The frames shown at higher magnification on right demonstrate the cell counting method, which incorporated two exclusion lines (left and bottom in red) and two inclusion lines (right and top in yellow) in each of the frames. A cell was counted if it showed clear nuclear staining, and was completely inside the sampling frame, or if it crossed the inclusion lines. Each counted cell has been marked with a colored line. The black arrows point to the estimated upper limit of layer 2, and lower limit of layer 6 in both sections. Neurons located below the estimated lower limits of layer 6 were regarded as white matter neurons, and not quantified in the present study.

Neurons were identified within each frame using previously published criteria (Schmitz and Hof 2005; Dorph-Petersen et al. 2009; Atapour et al. 2017). The cell counting method incorporated two exclusion lines (left and bottom) and two inclusion lines (right and top) in each of the counting frames (Fig. 1). A cell was counted if it was completely inside the frame or if it crossed the inclusion lines. Only cells with clear nuclear staining, independent of size and shape, were counted (the two counting strips illustrated in Figure 1 correspond to the red and green profiles shown in Supplementary Figure S1, for areas V1 and 8aV respectively). For each area, counting was performed by 2 of the present authors, each responsible for 2 strips. Given the high contrast of the NeuN stain, significant discrepancies between estimates obtained by different individuals were very rare, and in most cases could be explained by factors other than human error (e.g. different degrees of cortical curvature, or small folds in the sections). On the few occasions when human error was detected, an adjacent strip was selected, and new estimates obtained. The differences between original and recounted estimates were always <5%. All measurements were corrected using a shrinkage factor of 0.801 (by volume), which was obtained in an earlier study that used the same protocol (Atapour et al. 2017), based on measurements of the known distance between electrodes placed *in vivo*, and the separation between their tracks, visualized in NeuN-stained sections.

Average estimates of neuronal density were calculated for each area as the average of the estimates obtained in each of the 4 sampled sections. Each of these estimates included all frames between the top of layer 2 and layer 6, but excluded frames that had centers located in the white matter. For frames that straddled the boundary between layer 6 and the white matter, the percentage included in cortex was estimated, and this was taken into account in calculations of mean neuronal density. Estimates of neuronal density in layer 1 were calculated separately, and combined with those obtained for layers 2-6 for the calculation of number of neurons per unit column (see below).

### Additional analyses of neuronal density

To address the comparison of the present estimates of neuronal density with those reported by previous studies, we have performed additional analyses whereby distinct estimates of neuronal density were obtained from the same cortical locations of animal CJ167. These measurements were performed in 3 cortical areas characterized by very different laminar patterns: V1 (koniocortex), posterior parietal area PE (isocortex with 6 well defined layers), and the limb representation of the primary motor cortex (area 4ab; agranular cortex).

In the first analysis, we repeated the procedure outlined above using smaller counting frames (41 × 41 μm, separated by 70 μm). These parameters were chosen so as to reflect those employed in an earlier study of rodent and primate cortex (Charvet et al. 2015). Overlapping counting strips, from the same NeuN-stained sections, were used. The results, shown in Supplementary Figure S2, show that estimates of mean neuronal are not severely affected by the choice of frame size. Although local estimates fluctuated more noticeably when small counting frames are used, the global estimates of mean neuronal density obtained with different frame sizes only differed by 4.4-6.3%.

In the second analysis, we compared neuronal density estimates obtained in adjacent sections stained with the NeuN and Nissl stains. In counting Nissl-stained neurons we attempted to follow the same criteria employed by Cahalane et al. (2012) and Charvet et al. (2015): namely, “neurons were distinguished by the presence of coarse and dark stained Nissl substance in the cytoplasm, a distinct nucleus and a pale soma, contrasting with glial cells, which lack a conspicuous nucleolus and endoplasmic reticulum and exhibit a condensed soma”. This analysis, shown in Supplementary Figure S3, confirms that the estimates obtained in Nissl-stained sections are overall higher than those obtained with NeuN (Hilgetag et al. 2016), but also shows that estimates obtained in adjacent sections stained by the two methods are highly correlated (Gittins and Harrison 2004), revealing similar density profiles as a function of cortical depth. In general, the NeuN technique enables a clearer distinction between small neurons and glia.

### Estimating cortical thickness and number of neurons per unit column

Based on the neuronal density data we also obtained estimates of the number of neurons in columns extending from the surface of the brain to the bottom of layer 6. More specifically, we have estimated the number of neurons contained in a radially oriented compartment of cortex with 1 mm^2^ cross-section (parallel to the cortical layers). Estimates for each cytoarchitectural area were obtained using the formula:

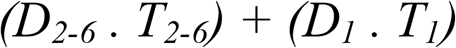

Where *D_2-6_* and *D_1_* are the mean neuronal densities in layers 2-6 and layer 1, respectively (in neurons/ mm^3^), and *T_2-6_* and *T_1_* are the thicknesses of layers 2-6 and layer 1 (in mm). To perform this calculation we estimated the thickness of the cortex as a whole, measured perpendicular to the cortical layers of the different areas, as well as estimates of the proportion of the thickness that was encompassed by layer 1.

To obtain the estimates of mean thickness of the cortex in the different areas, we employed a computational approach based on 3-d reconstruction and segmentation of the cortex. The calculations were performed using an in-house implementation of the two-surface Laplacian-based method (Jones et al., 2000; Waehnert et al. 2013) written in Python 2.7, and utilizing the Visualization Toolkit framework (Schroeder et al. 2006, http://www.vtk.org/). The process started with delineating the cortical hemisphere of CJ167 using the ITKSnap software (Yushkevich et al., 2006, http://itksnap.org). The pial surface and the white matter (WM) boundary were manually outlined (as shown in http://marmosetbrain.org). The contours were revised for artifacts which could affect the thickness estimates, such as connected banks of a sulcus. Each surface was then assigned with potential values (+1 for the WM boundary, -1 for the pial surface). Afterwards, a Laplacian equation was solved on a grid of one point per voxel, yielding the distribution of potential inside the cortical mask. A gradient of the potential was computed producing a dense (defined for each voxel) vector field. Sample points placed on the pial surface and propagated along the directions defined by the vector field were constrained to follow trajectories (streamlines) which 1) do not intersect with other trajectories or form loops, 2) cross the isosurfaces of the potential (including the pial surface and the WM boundary) at a right angle, and 3) generate an identical trajectory when propagated in the opposite direction. The length of each streamline provides an estimate of the thickness of the cortex at a specific point of the pial surface (Fig. 2).

**Figure 2.**
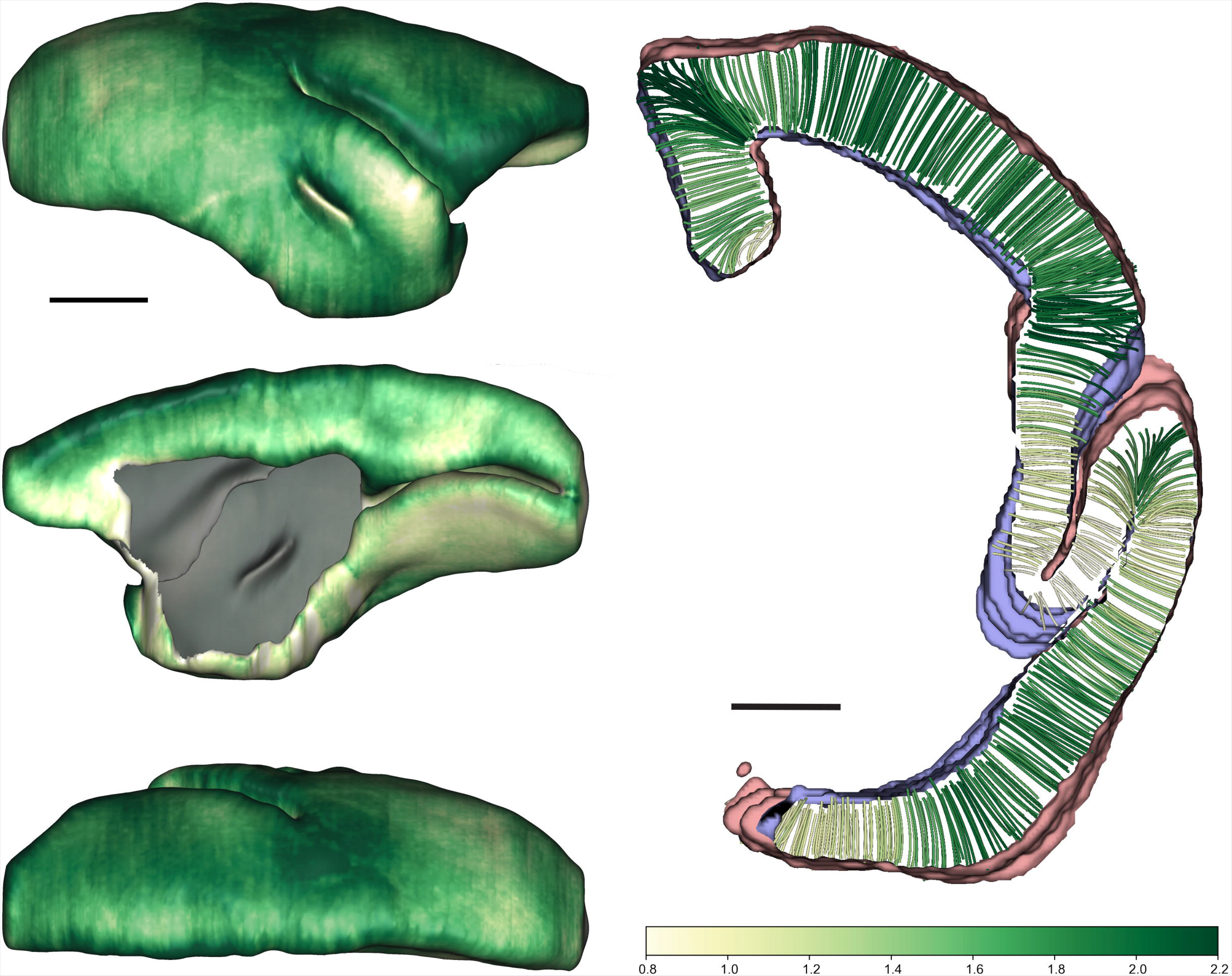
Determination of cortical thickness. **Left:** lateral, medial and dorsal views of the reconstructed cerebral cortex of animal CJ167. In this view, the thickness of the cortex (in mm) is indicated according to the color scale bar illustrated on the bottom right (also used in Fig. 10). Scale bar= 5 mm. **Right:** Method used to estimate thickness, based on the generation of streamlines perpendicular to isopotential surfaces in a two-surface Laplacian model (see “Materials and Methods” for details). This panel shows a 1 mm thick coronal slab at the level of the lateral sulcus in the right hemisphere. The generation of streamlines was performed in a 3-dimensional model, meaning that, at any given coronal level, many streamlines cross into adjacent sections. In this illustration, only streamlines which are fully enclosed within the 1 mm thick slab are shown, and among these only 1 in 20 (5%) are drawn, for clarity. Scale bar: 2 mm.

To reduce numerical errors, the outline of the cortex was smoothed with a Gaussian kernel (50×100×50 μm). Subsequently, over 376,000 points were defined on the pial surface (approximately 200 points per mm^2^, on average). Starting from each point a trajectory (streamline) was computed by traversing a distance of 20 μm per step, in a direction defined by the gradient vector. At each step, the corresponding cortical area was recorded. Upon reaching the WM boundary, the total length of the trajectory was calculated, and the streamline was assigned to the area to which the majority of the traversed locations was assigned. The lengths of all streamlines allotted to each area were averaged, yielding estimates of the mean thickness of the cortex in different areas (distributions of streamline lengths for several areas are shown in Supplementary Figure S4). The relative thickness of layer 1 (measured in sections) was then subtracted from this estimate, yielding the values of *T_1_* and T_2-6_ used for the calculation.

## Results

According to current knowledge, the marmoset cortex comprises 116 cytoarchitectural areas (Paxinos et al. 2012; Woodward et al. 2018), excluding the hippocampus (see Bunk et al. 2011 for estimates of neuronal density in the marmoset hippocampal formation) and the anterior olfactory nucleus. The areas included in the present study, and their abbreviations, are listed in Table 1. We identified each of the areas based primarily on cytoarchitectural features observable in NeuN-stained sections, but also using differences in myeloarchitecture and intensity of cytochrome staining as additional criteria to refine the delineations (e.g. Rosa et al. 2005, 2009; Palmer and Rosa 2006a, 2006b; Burman and Rosa 2009; Burman et al. 2006, 2014a, 2014b, 2015). We then used stereological techniques to obtain mean estimates of neuronal density for each area of animal CJ167 (Fig. 3, Table 1), as well as neuronal density profiles across the thickness of the cortex, from layer 2 to layer 6 (Fig. 4 and Supplementary Figure S1).

**Figure 3.**
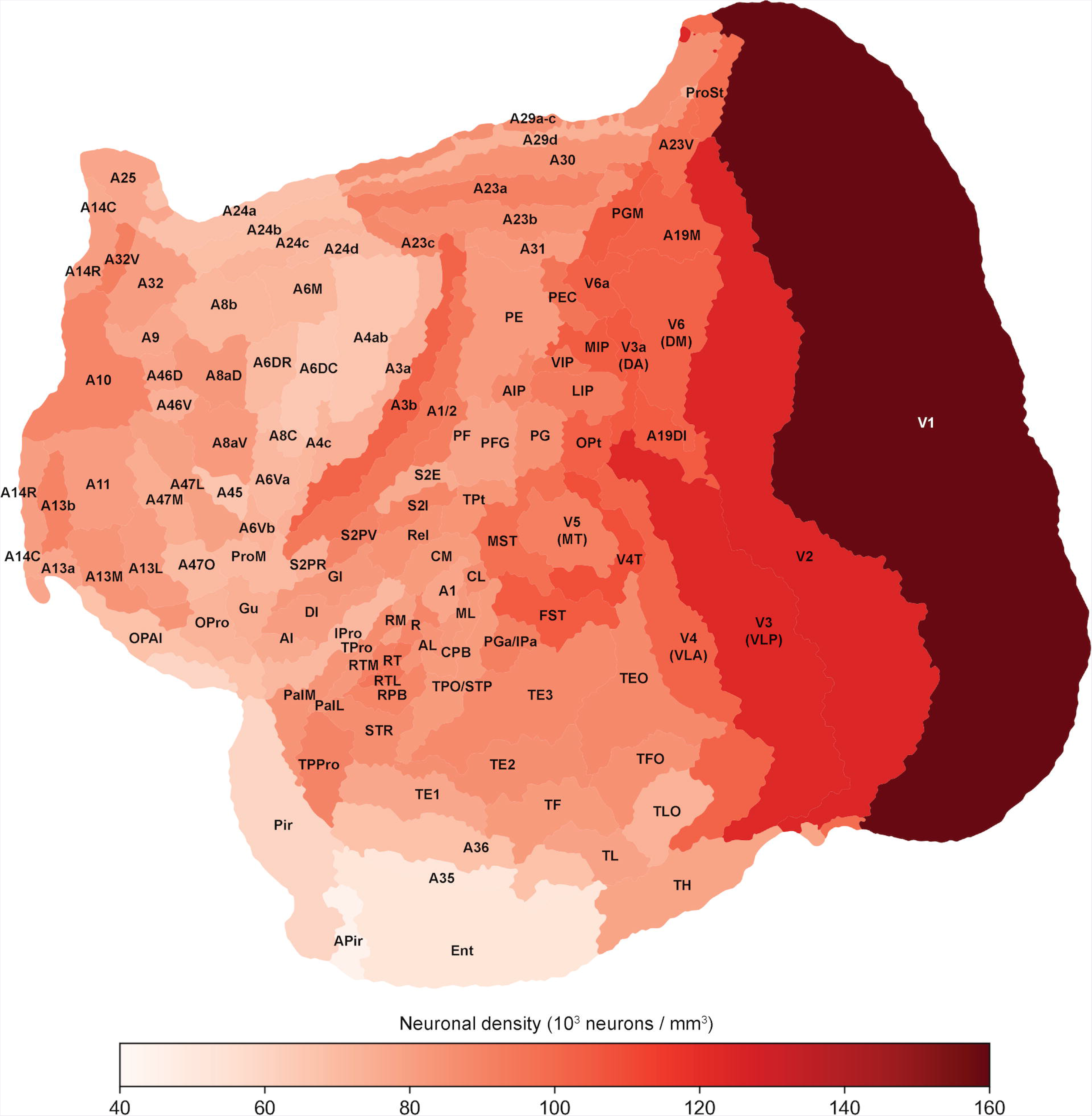
Estimates of mean neuronal density for 116 areas, shown in an “unfolded” representation of the marmoset cortex (Majka et al. 2016). Areal boundaries were estimated using the criteria illustrated by Paxinos et al. (2012). Abbreviations are listed in Table 1. To facilitate comparisons, in this and other figures, “flat maps” are represented according to the convention used by Paxinos et al. (2012) and adopted in the Marmoset Brain Architecture web site (www.marmosetbrain.org), i.e. anterior to the left, and medial to the top. A single scale bar cannot be given, in view of distortions inherent to forcing a 2-dimensional solution on a complex 3-dimensional surface. However, the total area represented here has been measured in the 3dimensional model as corresponding to 1095 mm^2^ at the level of the mid-thickness of the cortex.

**Figure 4.**
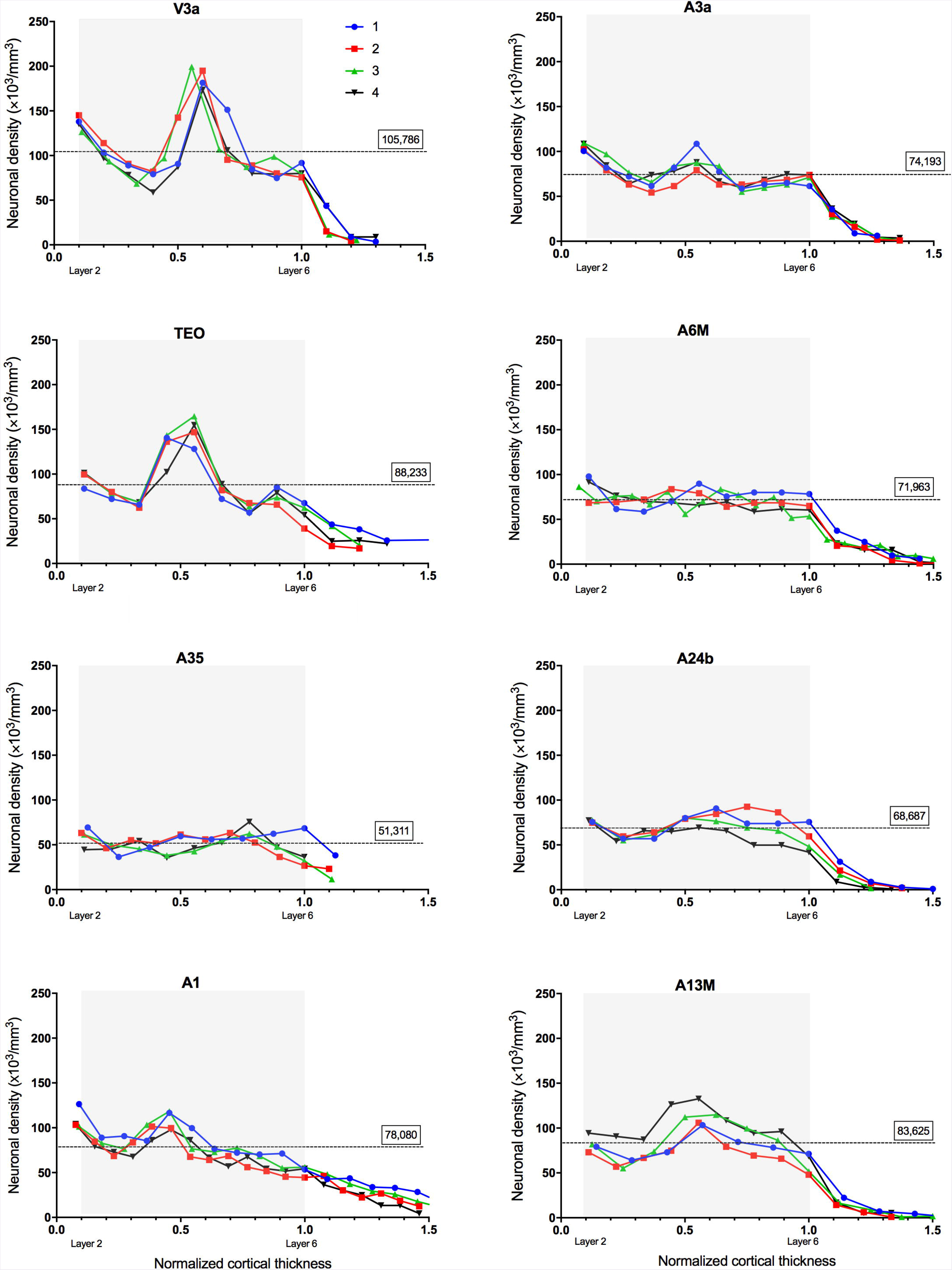
Variations of neuronal density across cortical depth for 8 areas of the marmoset cortex. Neuronal density calculated within each counting frame is plotted across the normalized depth of cortex (0= top of layer 2; 1.0= bottom of layer 6, estimated visually). In each graph, data from 4 columnar-shaped counting strips (shown in different colors) are presented. The gray shading represents our estimates of the cortex between the top of layer 2 and the bottom of layer 6. Values greater than 1 (outside gray box) represent density of white matter neurons. The dashed lines represent mean average densities, calculated as the mean of estimates obtained within the cortex in the 4 counting strips. Matching images of the 4 counting strips can be downloaded from http://www.marmosetbrain.org/cell_density. In this web site, the file name represents the section number, followed by the strip identifier (1-4). The same convention is used for all graphs (i.e. sample 1 in blue, sample 2 in red, sample 3 in green, and sample 4 in black). Values greater than 1 (outside gray box) represent density of white matter neurons. For abbreviations, see Table 1.

**Table 1:** Average neuronal densities in areas of the marmoset cortex

| Cortical area* |  | Neuronal density<br>(neurons/mm<sup>3</sup>) | Standard deviation<br>(neurons/mm<sup>3</sup>) |
| --- | --- | --- | --- |
| **Dorsolateral prefrontal cortex** |  |  |  |
| A10 | Area 10 | 89,069 | 17,886 |
| A9 | Area 9 | 77,175 | 16,746 |
| A46V | Area 46, ventral part | 74,153 | 16,328 |
| A46D | Area 46, dorsal part | 77,772 | 18,146 |
| A8aD | Area 8a, dorsal part | 82,351 | 15,910 |
| A8b | Area 8b | 72,198 | 13,409 |
| A8aV | Area 8a, ventral part | 81,908 | 18,716 |
| **Ventrolateral prefrontal cortex** |  |  |  |
| A47L | Area 47 (12), lateral part | 79,718 | 16,629 |
| A47M | Area 47 (12), medial part | 75,580 | 13,499 |
| A45 | Area 45 | 66,485 | 14,561 |
| A47O | Area 47 (12), orbital part | 70,445 | 15,576 |
| ProM | Proisocortical motor region (precentral opercular area) | 70,907 | 17,643 |
| **Orbitofrontal cortex** |  |  |  |
| A11 | Area 11 | 80,847 | 15,707 |
| A13b | Area 13b | 90,238 | 19,002 |
| A13a | Area 13a | 77,749 | 13,727 |
| A13L | Area 13, lateral part | 80,821 | 18,726 |
| A13M | Area 13, medial part | 83,625 | 20,974 |
| OPAI | Orbital periallocortex | 67,985 | 13,694 |
| OPro | Orbital proisocortex | 72,621 | 14,995 |
| Gu | Gustatory area | 74,314 | 18,211 |
| **Medial prefrontal cortex** |  |  |  |
| A32 | Area 32 | 80,761 | 14,809 |
| A32V | Area 32, ventral part | 91,321 | 14,539 |
| A14R | Area 14, rostral part | 80,346 | 15,996 |
| A14C | Area 14, caudal part | 76,515 | 15,652 |
| A25 | Area 25 | 77,930 | 10,628 |
| A24a | Area 24a | 68,238 | 14,896 |
| A24b | Area 24b | 68,687 | 12,737 |
| A24c | Area 24c | 72,652 | 9,731 |
| A24d | Area 24d | 70,959 | 10,272 |
| **Motor and premotor cortex** |  |  |  |
| A6DR | Area 6, dorsorostral part | 70,198 | 9,644 |
| A6Vb | Area 6, ventral, part b | 73,521 | 16,366 |
| A6Va | Area 6, ventral, part a | 71,164 | 13,770 |
| A8C | Area 8, caudal part | 68,530 | 13,097 |
| A6M | Area 6, medial (supplementary motor) part | 71,963 | 10,336 |
| A6DC | Area 6, dorsocaudal part | 64,917 | 14,026 |
| A4c | Area 4, part c (primary motor, representation of head musculature) | 72,041 | 13,400 |
| A4ab | Area 4, parts a and b (primary motor, representation of axial and limb musculature) | 65,959 | 12,690 |
| **Insula and rostral lateral sulcus cortex** |  |  |  |
| PaIM | Parainsular area, medial part | 83,808 | 19,854 |
| AI | Agranular insular area | 76,515 | 11,750 |
| PaIL | Parainsular area, lateral part | 82,547 | 19,667 |
| DI | Dysgranular insular area | 80,606 | 15,168 |
| GI | Granular insular area | 82,419 | 19,376 |
| Ipro | Insular proisocortex | 71,195 | 16,674 |
| Tpro | Temporal proisocortex | 87,064 | 16,577 |
| **Somatosensory cortex** |  |  |  |
| S2PR | Secondary somatosensory area, parietal rostral area | 74,173 | 20,732 |
| A3a | Area 3a | 74,193 | 14,786 |
| S2PV | Secondary somatosensory area, parietal ventral area | 89,991 | 24,852 |
| A3b | Area 3b (primary somatosensory area) | 101,536 | 26,105 |
| S2I | Secondary somatosensory area, internal part | 89,564 | 22,377 |
| S2E | Secondary somatosensory area, external part | 76,559 | 24,859 |
| A1/2 | Areas 1 and 2 | 94,083 | 27,727 |
| **Auditory cortex** |  |  |  |
| RTL | Rostrotemporal lateral area | 94,423 | 21,739 |
| RT | Rostrotemporal area | 93,525 | 20,457 |
| RPB | Rostral parabelt area | 89,296 | 19,420 |
| RTM | Rostrotemporal medial area | 91,076 | 16,830 |
| R | Rostral auditory area | 89,089 | 23,058 |
| RM | Rostromedial area | 79,803 | 17,763 |
| AL | Anterolateral area | 83,909 | 20,668 |
| A1 | Primary auditory area | 78,080 | 19,981 |
| CM | Caudomedial area | 81,552 | 18,616 |
| CPB | Caudal parabelt area | 81,425 | 21,249 |
| ML | Middle lateral area | 82,588 | 20,734 |
| CL | Caudolateral area | 86,610 | 22,009 |
| **Lateral and inferior temporal cortex** |  |  |  |
| TPPro | Temporopolar proisocortex | 90,130 | 17,549 |
| STR | Superior temporal rostral area | 86,521 | 17,753 |
| TE1 | Temporal area TE, part 1 | 77,335 | 24,078 |
| TPO/STP | Temporo-parieto-occipital association area | 81,476 | 24,209 |
| ReI | Retroinsular area | 84,220 | 21,635 |
| TE2 | Temporal area TE, part 2 | 85,907 | 32,487 |
| PGa/IPa | Areas PGa and IPa (fundus of superior temporal ventral area) | 91,229 | 29,663 |
| TPt | Temporoparietal transitional area | 83,428 | 21,195 |
| TE3 | Temporal area TE, part 3 | 87,504 | 28,727 |
| TEO | Temporal area TE, occipital transition part | 88,233 | 31,768 |
| **Ventral temporal cortex** |  |  |  |
| Pir | Piriform area | 59,871 | 16,288 |
| APir | Amygdalopiriform transition area | 44,745 | 13,685 |
| Ent | Entorhinal area | 51,867 | 15,459 |
| A36 | Area 36 | 67,998 | 19,672 |
| A35 | Area 35 | 51,311 | 11,297 |
| TF | Temporal area TF | 81,484 | 27,093 |
| TL | Temporal area TL | 79,111 | 22,371 |
| TH | Temporal area TH | 78,537 | 21,606 |
| TLO | Temporal area TL, occipital transition part | 73,716 | 22,922 |
| TFO | Temporal area TF, occipital transition part | 85,358 | 26,976 |
| **Posterior cingulate, medial and retrosplenial cortex** |  |  |  |
| A23c | Area 23c | 87,820 | 21,512 |
| A23a | Area 23a | 90,649 | 20,537 |
| A29d | Area 29d | 74,446 | 13,941 |
| A30 | Area 30 | 84,089 | 15,441 |
| A23b | Area 23b | 85,066 | 22,365 |
| A29a-c | Area 29, parts a-c | 85,477 | 20,548 |
| A23V | Area 23, ventral part | 98,063 | 30,049 |
| ProSt | Prostriate area | 98,139 | 21,147 |
| **Posterior parietal cortex** |  |  |  |
| PF | Parietal area PF | 92,283 | 25,919 |
| PE | Parietal area PE | 82,143 | 21,120 |
| PFG | Parietal area PFG | 78,936 | 24,508 |
| A31 | Area 31 | 81,862 | 22,445 |
| AIP | Anterior intraparietal area | 87,971 | 28,927 |
| PG | Parietal area PG | 85,114 | 27,985 |
| PEC | Parietal area PE, caudal transitional part | 95,478 | 29,458 |
| VIP | Ventral intraparietal area | 93,539 | 29,939 |
| LIP | Lateral intraparietal area | 93,038 | 30,399 |
| PGM | Parietal area PG, medial part | 104,582 | 30,089 |
| V6a | Visual area 6a (posterior parietal medial area) | 102,372 | 33,052 |
| OPt | Occipito-parietal transitional area | 102,916 | 37,958 |
| MIP | Medial intraparietal area | 104,502 | 34,330 |
| **Visual cortex** |  |  |  |
| MST | Medial superior temporal area | 97,815 | 33,888 |
| FST | Fundus of superior temporal sulcus area | 104,832 | 32,927 |
| V5 | Visual area 5 (middle temporal area, MT) | 91,754 | 27,866 |
| V4T | Visual area 4, transitional part (middle temporal crescent) | 109,315 | 38,132 |
| A19M | Area 19, medial part | 102,466 | 35,991 |
| V3a | Visual area 3a (dorsoanterior area, DA) | 105,786 | 35,580 |
| V4 | Visual area 4 (ventrolateral anterior area, VLA) | 101,872 | 36,046 |
| V6 | Visual area 6 (dorsomedial area, DM) | 102,426 | 34,077 |
| A19DI | Area 19, dorsointermediate part (DI) | 104,911 | 32,325 |
| V3 | Visual area 3 (ventrolateral posterior area, VLP) | 123,176 | 39,658 |
| V2 | Visual area 2 | 124,091 | 41,392 |
| V1 | Primary visual area | 159,450 | 34,737 |

\* Designations of cortical areas according to Paxinos et al. 2012. Within each broad subdivision of cortex, areas are listed in rostral to caudal order.

In agreement with previous studies, we found significant variations in neuronal density among cytoarchitectural areas (Figs. 3-6, and Table 1). In the marmoset, estimates encompassing layers 2 to 6 cover an approximately threefold range, with V1 showing the highest density (>150,000 neurons/ mm^3^; e.g. Fig. 1), and allocortical areas such as piriform, entorhinal and perirhinal areas exhibiting the lowest (~50,000 neurons/ mm^3^; e.g. Fig. 5, area 35). As detailed below, low densities (~68,000 neurons/ mm^3^) were characteristic of many proisocortical and periallocortical areas, including, for example, ventral subdivisions of the anterior cingulate cortex (e.g. areas 24a and 24b), the orbital proisocortex (OPro) and periallocortex (OPAl), and the perirhinal area (area 36). In addition, subdivisions of the motor and premotor complex (Bakola et al. 2015) were characterized by relatively low densities (64,000 – 73,000 neurons/ mm^3^) in comparison with adjacent areas. Overall, the data confirmed the existence of a coarse posterior to anterior gradient, with neuronal density trending from high to low values along this axis; this was apparent whether the analysis included all cortical areas (Fig. 6) or only isocortical areas (as in Cahalane et al. 2012; Supplementary Figure S5; Supplementary Table S1). However, as detailed below, there is significant granularity in this gradient, which leads to local deviations from a strict posterior to anterior order in many regions of the cortex.

**Figure 5.**
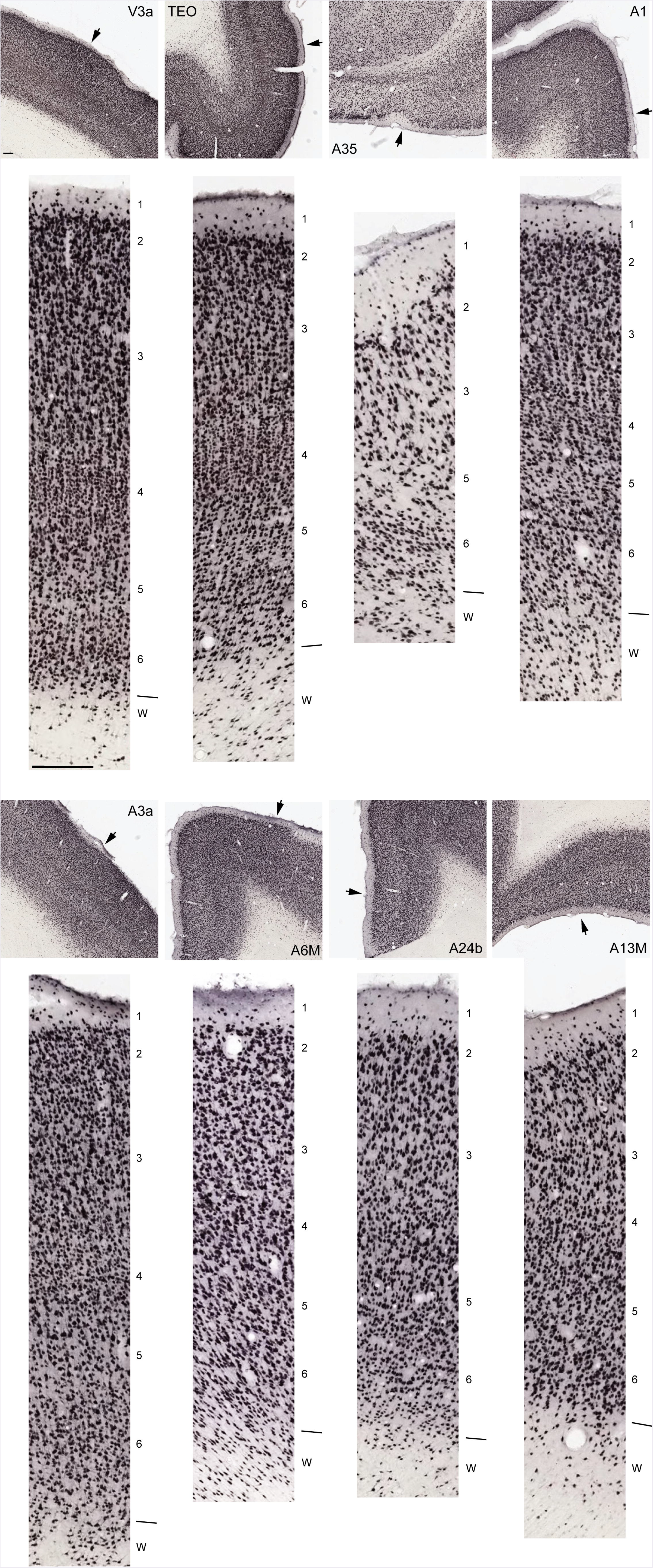
Examples of NeuN staining of the areas for which neuronal density profiles are illustrated in Figure 4. The arrows in the upper panels point to the locations from which the high magnification images were obtained. The approximate levels of the centers of cortical layers and the boundary between cortex and white matter (W) are identified on each high-resolution image. Scale bars (shown in the upper and lower panels for area V3a, top left) equal 100 μm.

**Figure 6:**
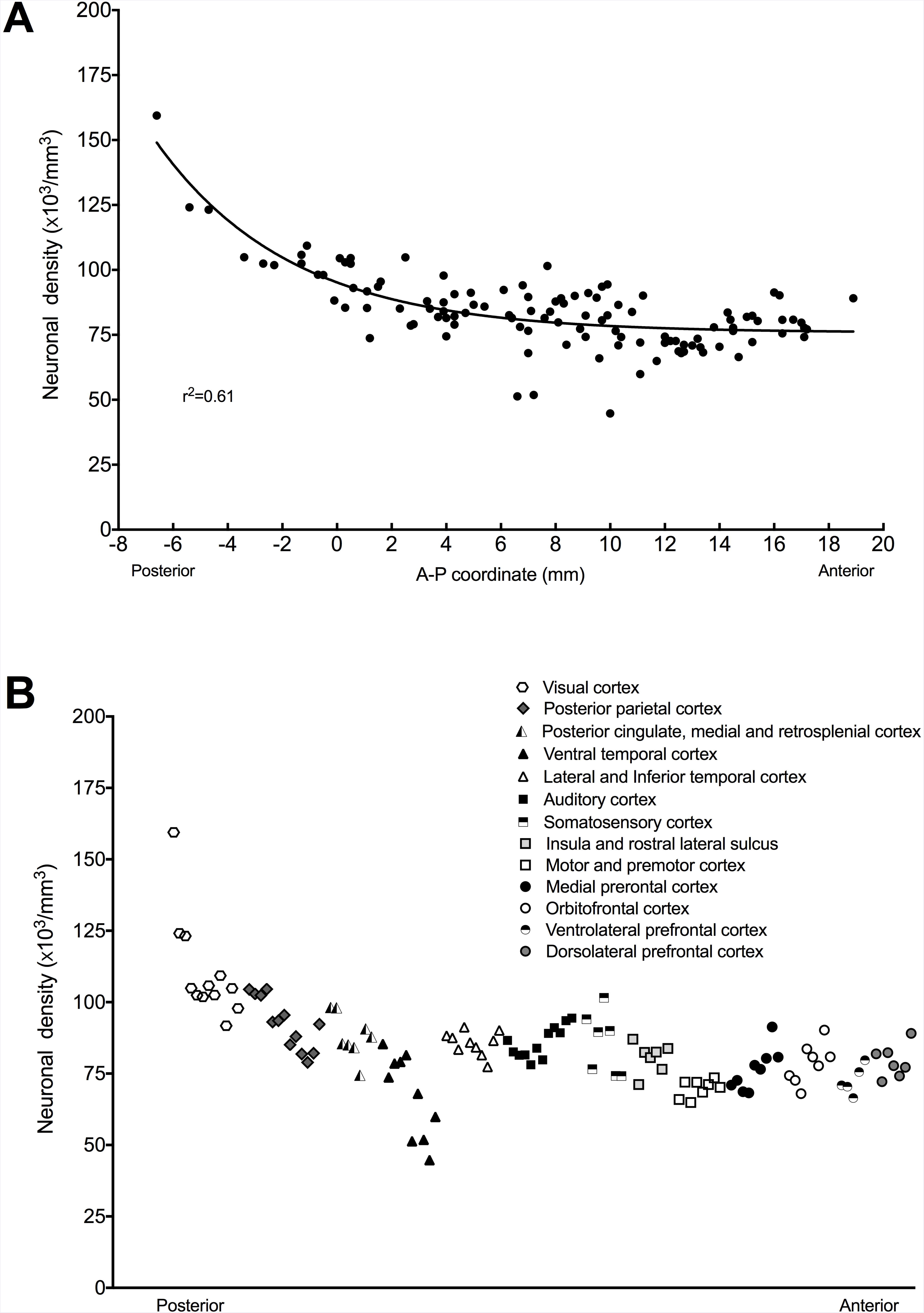
**A-** Neuronal density as a function of anteroposterior (A-P) coordinate. The A-P coordinate of each area is assigned according to its barycenter, calculated in a 3-dimensional reconstruction aligned to a marmoset brain template using the procedure described by Majka et al. (2016). The line represents a fitted exponential function, which provides an resonable (r^2^=0.61) fit to the data. **B-** Same data, with the areas grouped according to the functional groups given in Table 1. In this representation, the groups of areas are ordered according to the mean A-P coordinate of the barycenters of areas; within each group, areas are placed in order of A-P coordinate.

Neuronal density estimates for layer 1 are presented in Table 2. Although the values obtained for different areas fluctuated, even within broad functional groups, this analysis did not reveal any clear gradients or trends (Supplementary Figure S6). We did observe differences in the distribution of white matter neurons, particularly with respect to how far these extended below layer 6 (e.g. Figs. 1, 4, 5, and Supplementary Figure S1), but these were not easily interpretable in the context of the present report. In particular, the lack of cytoarchitectural criteria made it impossible to assign these cells to specific areas, and it remains unclear how they are functionally related to the overlying areas. Hence, these neurons were not subject to quantification in the present study. In the Results presented below, unless otherwise specified, references to “neuronal density” correspond to density measured from the top of layer 2 to the estimated bottom of layer 6.

**Table 2:**
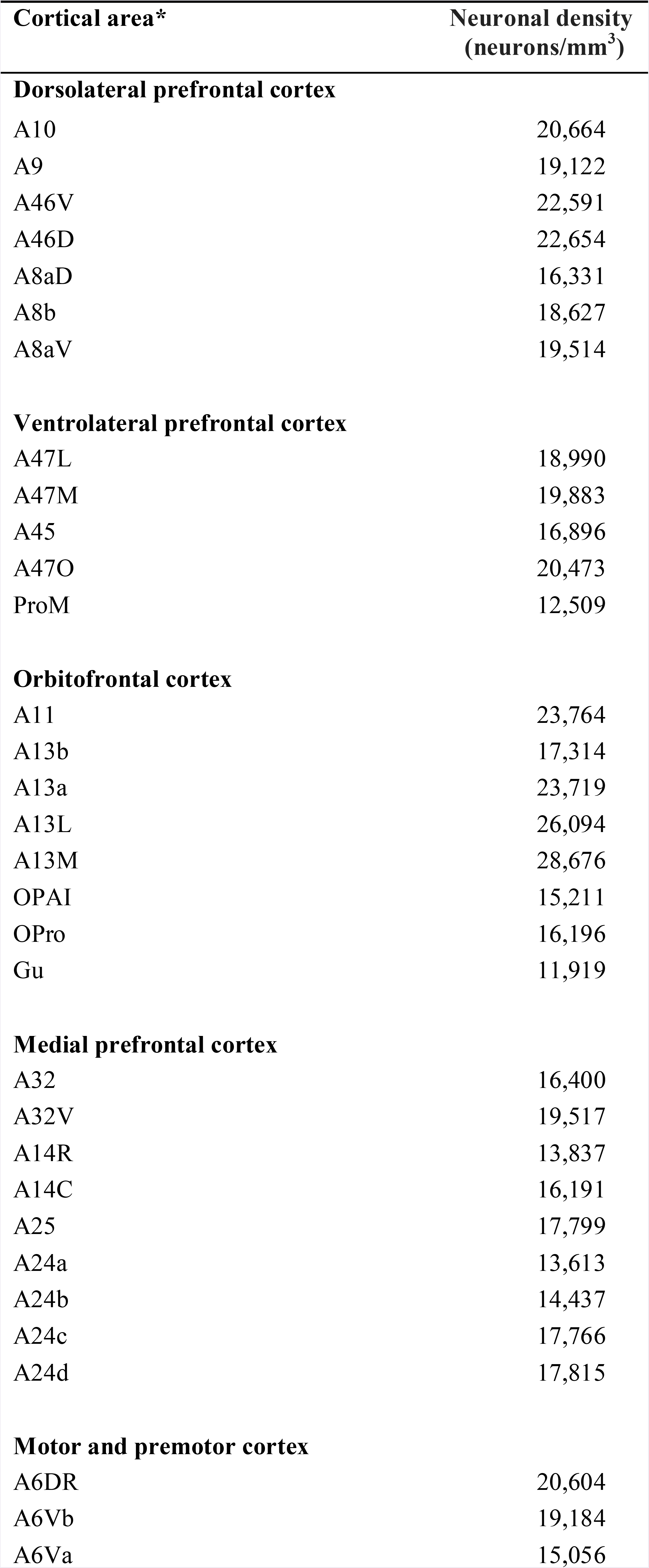
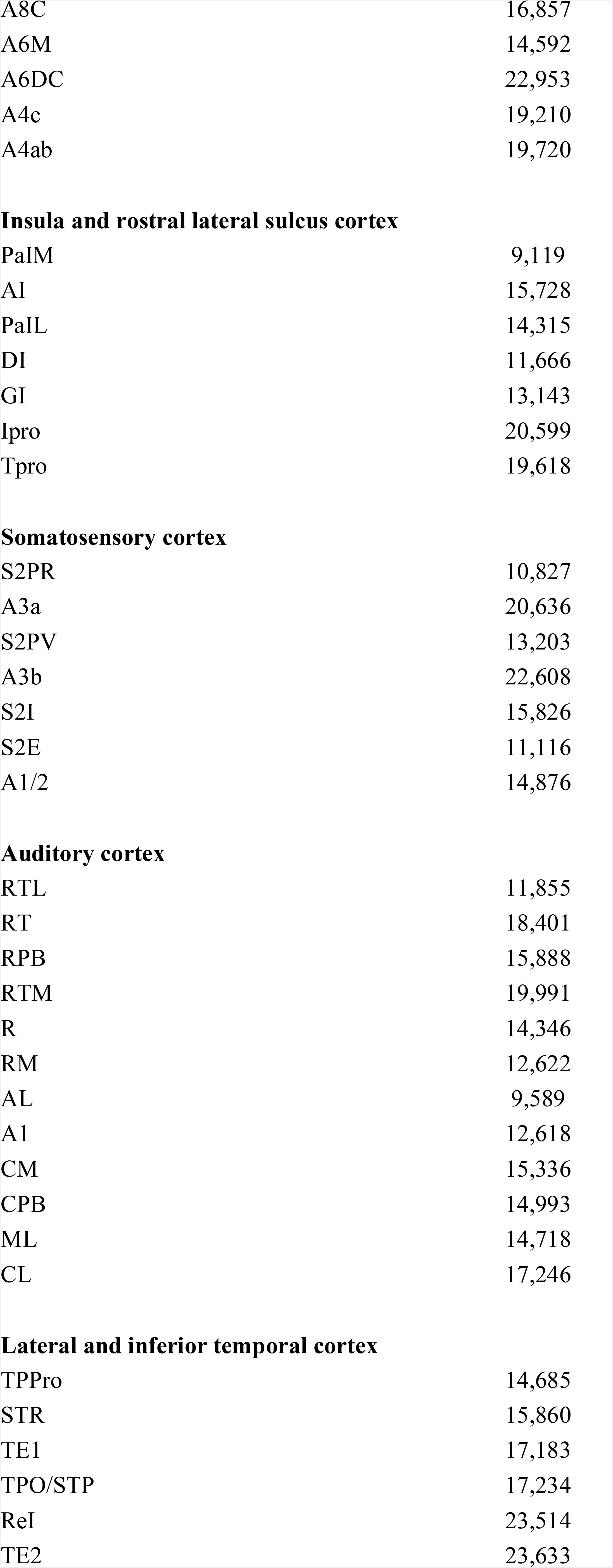
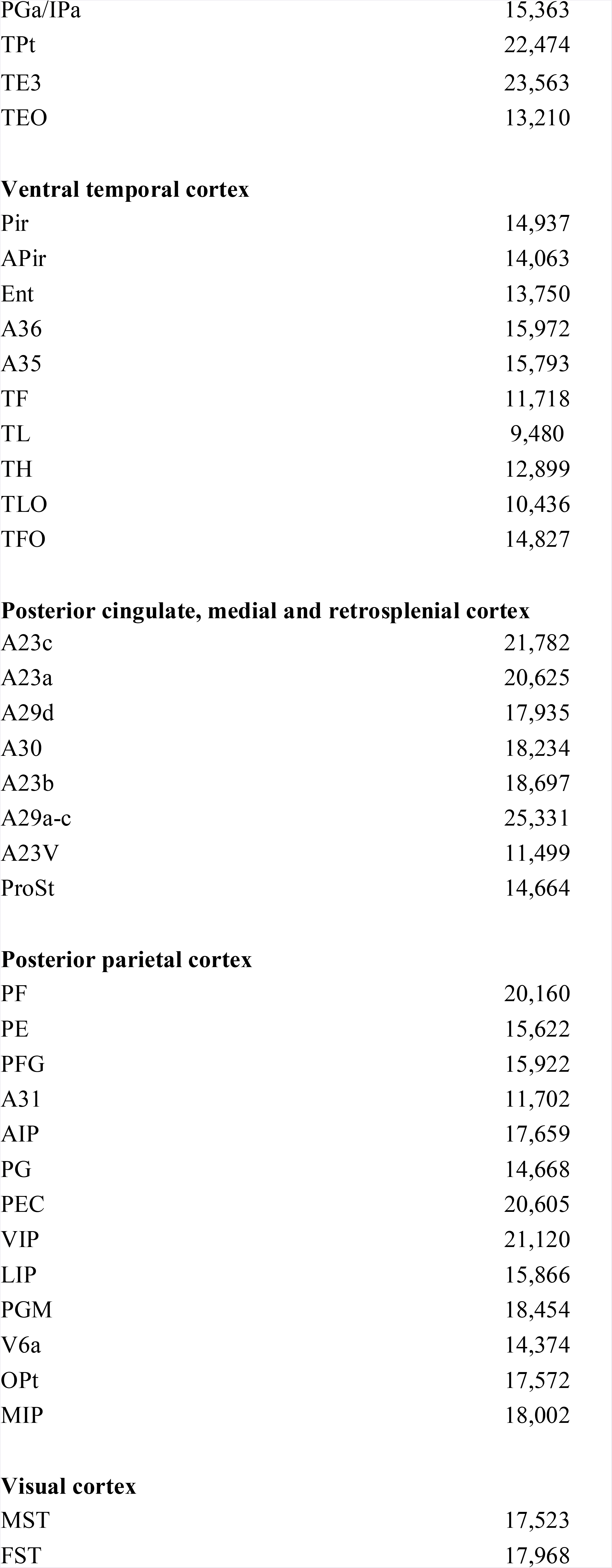
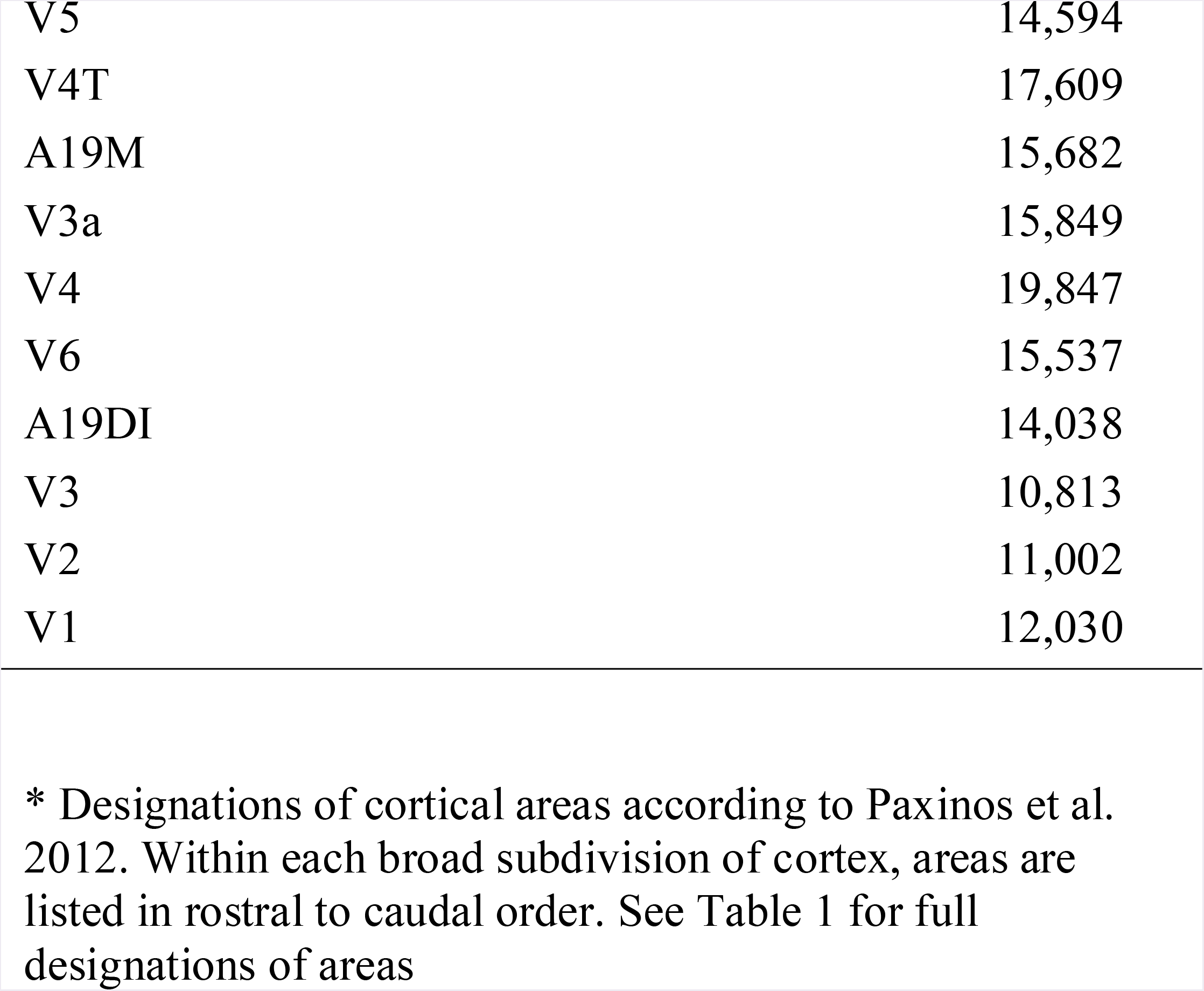
Average neuronal densities in layer 1 in areas of the marmoset cortex

| **Cortical area\*** | **Neuronal density**<br>**(neurons/mm<sup>3</sup>)** |
| --- | --- |
| **Dorsolateral prefrontal cortex** |  |
| A10 | 20,664 |
| A9 | 19,122 |
| A46V | 22,591 |
| A46D | 22,654 |
| A8aD | 16,331 |
| A8b | 18,627 |
| A8aV | 19,514 |
| **Ventrolateral prefrontal cortex** |  |
| A47L | 18,990 |
| A47M | 19,883 |
| A45 | 16,896 |
| A47O | 20,473 |
| ProM | 12,509 |
| **Orbitofrontal cortex** |  |
| A11 | 23,764 |
| A13b | 17,314 |
| A13a | 23,719 |
| A13L | 26,094 |
| A13M | 28,676 |
| OPAI | 15,211 |
| OPro | 16,196 |
| Gu | 11,919 |
| **Medial prefrontal cortex** |  |
| A32 | 16,400 |
| A32V | 19,517 |
| A14R | 13,837 |
| A14C | 16,191 |
| A25 | 17,799 |
| A24a | 13,613 |
| A24b | 14,437 |
| A24c | 17,766 |
| A24d | 17,815 |
| **Motor and premotor cortex** |  |
| A6DR | 20,604 |
| A6Vb | 19,184 |
| A6Va | 15,056 |
| A8C | 16,857 |
| A6M | 14,592 |
| A6DC | 22,953 |
| A4c | 19,210 |
| A4ab | 19,720 |
| **Insula and rostral lateral sulcus cortex** |  |
| PaIM | 9,119 |
| AI | 15,728 |
| PaIL | 14,315 |
| DI | 11,666 |
| GI | 13,143 |
| Ipro | 20,599 |
| Tpro | 19,618 |
| **Somatosensory cortex** |  |
| S2PR | 10,827 |
| A3a | 20,636 |
| S2PV | 13,203 |
| A3b | 22,608 |
| S2I | 15,826 |
| S2E | 11,116 |
| A1/2 | 14,876 |
| **Auditory cortex** |  |
| RTL | 11,855 |
| RT | 18,401 |
| RPB | 15,888 |
| RTM | 19,991 |
| R | 14,346 |
| RM | 12,622 |
| AL | 9,589 |
| A1 | 12,618 |
| CM | 15,336 |
| CPB | 14,993 |
| ML | 14,718 |
| CL | 17,246 |
| **Lateral and inferior temporal cortex** |  |
| TPPro | 14,685 |
| STR | 15,860 |
| TE1 | 17,183 |
| TPO/STP | 17,234 |
| ReI | 23,514 |
| TE2 | 23,633 |
| PGa/IPa | 15,363 |
| TPt | 22,474 |
| TE3 | 23,563 |
| TEO | 13,210 |
| **Ventral temporal cortex** |  |
| Pir | 14,937 |
| APir | 14,063 |
| Ent | 13,750 |
| A36 | 15,972 |
| A35 | 15,793 |
| TF | 11,718 |
| TL | 9,480 |
| TH | 12,899 |
| TLO | 10,436 |
| TFO | 14,827 |
| **Posterior cingulate, medial and retrosplenial cortex** |  |
| A23c | 21,782 |
| A23a | 20,625 |
| A29d | 17,935 |
| A30 | 18,234 |
| A23b | 18,697 |
| A29a-c | 25,331 |
| A23V | 11,499 |
| ProSt | 14,664 |
| **Posterior parietal cortex** |  |
| PF | 20,160 |
| PE | 15,622 |
| PFG | 15,922 |
| A31 | 11,702 |
| AIP | 17,659 |
| PG | 14,668 |
| PEC | 20,605 |
| VIP | 21,120 |
| LIP | 15,866 |
| PGM | 18,454 |
| V6a | 14,374 |
| OPt | 17,572 |
| MIP | 18,002 |
| **Visual cortex** |  |
| MST | 17,523 |
| FST | 17,968 |
| V5 | 14,594 |
| V4T | 17,609 |
| A19M | 15,682 |
| V3a | 15,849 |
| V4 | 19,847 |
| V6 | 15,537 |
| A19DI | 14,038 |
| V3 | 10,813 |
| V2 | 11,002 |
| V1 | 12,030 |

\* Designations of cortical areas according to Paxinos et al. 2012. Within each broad subdivision of cortex, areas are listed in rostral to caudal order. See Table 1 for full designations of areas

In order to ascertain the extent to which the differences observed in case CJ167 are representative of general trends, we also analyzed a sample of areas with different neuronal densities, using identical techniques, in two additional animals (cases CJ173 and CJ800) of different sex (males). This analysis, illustrated in Figure 7, shows that the estimates obtained in CJ167 are well in line with those obtained in the other cases, and that the inter-individual variability in neuronal density values is relatively small in comparison with the differences between areas, in animals of comparable age.

**Figure 7.**
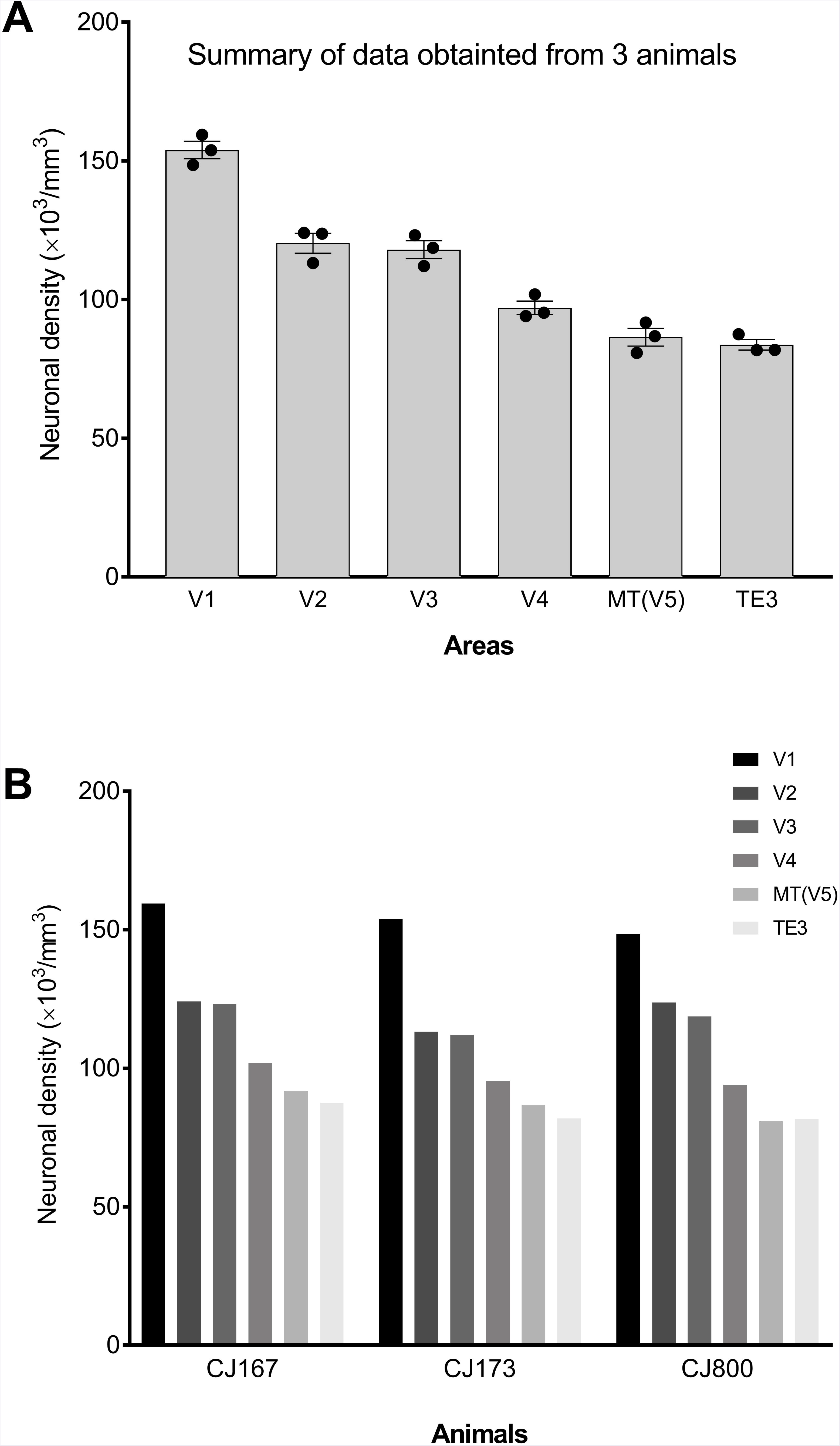
Analysis of inter-individual variability, based on a sample of 6 visual areas studied with identical methods in 3 animals. **A:** mean neuronal density. The individual circles indicate values obtained in 3 individuals. **B:** Same data, sorted by animal. Consistent trends are observed.

### Visual cortex

The data revealed a relatively ordered decrease in neuronal density along the posterior to anterior axis of the visual cortex (Figs. 3, 7). For example, after the exceptionally high neuronal density in V1 (159,450 neurons/ mm^3^), the second highest neuronal densities were found in visual areas 2 (V2; 124,091 neurons/ mm^3^) and 3 (V3, defined here as corresponding to the ventrolateral posterior area, VLP; see Angelucci and Rosa 2015; 123,176 neurons/ mm^3^). Notably, the dorsal component of the third visual complex (dorsomedial area, DM, or V6) had a much lower neuronal density (102,426 neurons/ mm^3^) than V3/ VLP, confirming the existence of significant asymmetries in the cortex rostral to V2 (Rosa and Tweedale 2000; Rosa et al. 2005; Jeffs et al. 2013, 2015). All other topographically organized extrastriate areas showed relatively high neuronal densities, around 100,000/mm^3^, although the middle temporal area (MT, or V5) represented an outlier, with a relatively lower neuronal density (91,754 neurons/ mm^3^), which stood out even in comparison with the adjacent medial superior temporal (MST) and fundus of superior temporal (FST) areas (97,815 neurons/ mm^3^ and 104,832 neurons/mm^3^, respectively). In the inferior temporal cortex visual association cortex, neuronal densities decreased from posterior (TEO: 88,233 neurons/ mm^3^) to anterior (TE1: 77,335 neurons/ mm^3^) areas. A similar anteroposterior trend was observed among visual association areas of the posterior parietal cortex, and the frontal areas most closely associated with the visual system (cytoarchitectural areas 8aV and 45; Burman et al. 2006; Reser et al. 2013) showed even lower neuronal densities (see below).

Given that some visual areas are particularly large, it is appropriate to ask if anteroposterior gradients of neuronal density can also be observed *within* these areas. In order to address this, we obtained additional measurements in some of the large visual areas, in order to extend the anteroposterior range of the samples. Supplementary Figure S7 (A, B) shows the results for areas V1 (the largest subdivision of the marmoset cortex) and TE3 (a subdivision of inferior temporal cortex which spans a large anteroposterior extent on the lateral surface of the temporal lobe). This analysis revealed that there is no systematic variation in neuronal density according to rostrocaudal level within V1. In TE3, for which the estimation of cytoarchitectural boundaries is subject to a greater error, the same analysis revealed somewhat lower neuronal density near its anterior one-third, compared to the posterior two-thirds (78,540 neurons/ mm^3^ versus 87,504 neurons/ mm^3^). The potential implications of such differences, which may suggest subdivisions within what are currently recognized as single, large areas, are discussed below.

### Somatosensory cortex

The highest neuronal density among somatosensory areas was found in cytoarchitectural area 3b (101,536 neurons/ mm^3^; Fig. 3), which corresponds to the somatosensory koniocortex (Krubitzer and Kaas 1990). An anterior to posterior gradient of decreasing neuronal density characterized the transitions to area 1/2 (a single cytoarchitectural field that in marmosets occupies the expected locations of areas 1 and 2; Paxinos et al. 2012; 94,083 neurons/ mm^3^) and to parietal somatic association areas PE (also known as area 5; Padberg et al. 2007; 82,143 neurons/ mm^3^) and PF (92,283 neurons/ mm^3^). Rostrally, area 3a, which has functional and anatomical characteristics that indicate both somatosensory and motor functions (Huffman and Krubitzer 2001), had a lower value of neuronal density (74,193 neurons/ mm^3^), which was intermediate between those found in the somatosensory and motor areas (see below). Among the secondary somatosensory complex areas, the lowest neuronal density was found anteriorly, in the parietal rostral area (S2PR; 74,173 neurons/ mm^3^). There was a significant difference between the external (S2E) and internal (S2I) cytoarchitectural segments of the secondary somatosensory area, which are located in the shoulder and medial bank of the lateral sulcus, respectively; this could be a result of different neuronal arrangements imposed by developmental mechanisms in regions with high and low cortical convexity (Hilgetag and Barbas 2006).

### Auditory cortex

Differences between auditory areas were relatively modest. Unlike among visual and somatosensory areas, there was little indication of a clear gradient of neuronal density that could be related to hierarchical levels of processing (Kaas and Hackett 2000): the auditory core, belt and parabelt areas had values of neuronal density that varied within similar ranges. However, within each of these broad functional subdivisions, there was a trend for highest densities to occur at rostral levels (Figs. 3, 6B). For example, the neuronal density in the rostrotemporal core area (RT; 93,525 neurons/ mm^3^) was higher than in the primary auditory area (A1; 78,080 neurons/ mm^3^), and that in the rostral parabelt (RPB; 89,296 neurons/ mm^3^) was higher than in the caudal parabelt (CPB; 81,425 neurons/ mm^3^). Likewise, among the belt areas, the highest density was found in the rostrotemporal lateral area (RTL; 94,423 neurons/ mm^3^). As shown below, differences in neuronal density among core areas were in part compensated by variations in the thickness of the cortex, resulting in a different spatial gradient when the number of neurons per column was calculated for these areas.

### Motor and premotor cortex

The marmoset primary motor cortex has different cytoarchitectural characteristics in the face/ oral cavity (area 4c) and body (areas 4a/b) representations, with large layer 5 pyramidal neurons (Betz cells) being concentrated in the latter subdivisions (Burman et al. 2008, 2014a). This qualitative difference was borne out in our quantitative analysis: cytoarchitectural area 4a/b had a lower neuronal density than area 4c (65,959 neurons/ mm^3^ vs. 72,041 neurons/ mm^3^), and within area 4a/b relatively low neuronal densities were observed near the midline (Supplementary Figure 7). In general, differences among motor and premotor areas were modest, and did not seem to conform to a spatial gradient; rather these areas of the caudal frontal lobe combined to form a region of relatively low neuronal density in comparison with the somatosensory cortex that lies at its caudal border, and the dorsolateral prefrontal cortex that lies at its rostral border (Figs. 3, 6B; see also Young et al. 2013; Turner et al. 2016).

### Prefrontal cortex

As summarized in Figure 3, estimates of neuronal density among areas in the dorsolateral, ventrolateral, medial and orbitofrontal subdivisions of the prefrontal cortex tended to be lower than those in the occipital and parietal lobes, but higher than those in motor and premotor areas. Remarkably, some of the highest values among prefrontal areas were in fact observed near the frontal pole, including area 10 (89,069 neurons/ mm^3^), area 13b (90,238 neurons/ mm^3^) and area 32V (91,321 neurons/ mm^3^; also known as area 10mc; Saleem et al. 2008; Burman and Rosa 2009). Conversely, throughout the prefrontal eulaminate cortex, the lowest values tended to occur caudally (e.g. area 8b, 72,198 neurons/ mm^3^; area 45, 66,485neurons/ mm^3^; area 47O, 70,445 neurons/ mm^3^). Low densities were also observed in the orbital proisocortical (OPro) and periallocortical (OPAl) areas (72,621 and 67,985 neurons/ mm^3^, respectively), in the proisocortical motor area (ProM; 70,907 neurons/ mm^3^), and in the gustatory cortex (Gu), which spans parts of the caudal lateral orbital surface and anterior insula (74,314 neurons/ mm^3^).

### Temporal association cortex and insula

Areas of the superior temporal association cortex, which are likely to have polysensory integration roles (Baylis et al. 1987; Rosa et al. 2018), had neuronal densities within a similar range to that observed in the adjacent inferior temporal visual association cortex. The maximum value (91,229 neurons/ mm^3^) was observed in cytoarchitectural area PGa/IPa, which follows the fundus of the superior temporal sulcus, leading to marked compression of the cortex. Among other subdivisions, values ranged between 81,476 neurons/ mm^3^ (cytoarchitectural area TPO) and 90,130 neurons/ mm^3^ (temporal pole proisocortex [TPPro]. This range also encompasses the neuronal densities observed in lateral sulcus areas such as those in the parainsular complex (~83,000 neurons/ mm^3^) and the temporal proisocortex (TPro; 87,064 neurons/ mm^3^). In the insular complex the neuronal densities tended to be lower, ranging from 71,195 neurons/ mm^3^ (insular proisocortex, Ipro) to 82,419 neurons/ mm^3^ (granular insular cortex, GI). In the ventral temporal (parahippocampal and perirhinal) areas, neuronal density values were also generally low, with a trend for increasing densities from ventromedial (e.g. area 36; 67,998 neurons/ mm^3^) to lateral (e.g. TFO; 85,358 neurons/ mm^3^) subdivisions – that is, as one considers areas progressively further away from the entorhinal and hippocampal complexes.

### Posterior parietal cortex

Relatively high neuronal densities were observed in visuomotor association areas of the posterior parietal cortex (Figs. 3, 6B). Caudal subdivisions involved in visuomotor integration, including V6a, the medial intraparietal area (MIP), and cytoarchitectural areas OPt and PGM tended to be characterized by higher neuronal densities (~100,000 neurons/ mm^3^) in comparison with rostral areas, such as the ventral (VIP) and lateral (LIP) intraparietal areas (VIP, LIP; ~93,000 neurons/ mm^3^), the anterior intraparietal area (AIP; 87,971 neurons/ mm^3^) and cytoarchitectural area PFG (78,936 neurons/ mm^3^). This trend continued further rostrally, with even lower values being observed in cytoarchitectural areas PE (82,143 neurons/ mm^3^) and 31 (81,862 neurons/ mm^3^), which have been implicated in somatomotor association function (Passarelli et al. 2018).

### Cingulate and retrosplenial cortex

Similar to the posterior parietal cortex, the highest neuronal density values in the posterior cingulate/ retrosplenial complex occurred caudally (area 23V, 98,063 neurons/ mm^3^; ProSt [area prostriata], 98,139 neurons/ mm^3^). Estimates of the neuronal density among the rostral subdivisions of area 23 (areas 23a-c) tended to be lower (85,066 – 90,649 neurons/ mm^3^), and those in the anterior cingulate and subgenual areas (areas 24a-d and 25) were even lower (68,238 – 77,930 neurons/ mm^3^).

### Differences through the depth of the cortex

Qualitative differences in the thickness and degree of definition of cortical layers, and in neuronal density within these layers, have been the basis of cytoarchitectural analyses for over a century. Thus, it is not surprising that our analysis revealed clear differences in neuronal density profiles through the depth of different areas. Some of the observed variation is illustrated by examples given in Figures 4 and 5 (the data for all 116 areas is presented in Supplementary Figure S1). For example, profiles for sensory areas such as V3a (dorsoanterior area; Rosa and Schmid 1995), TEO, and A1 revealed clear peaks around the cortical midthickness, corresponding to layer 4, although the relative height and location of this peak varied. Relatively high neuronal densities in layer 2 were clear in some areas (e.g. visual area V3a, A1, somatosensory area 3a) but less so in others. In the medial subdivision of area 6 (6M), which is illustrated here as an example of an area in the motor/premotor complex, this analysis revealed comparatively little variation in neuronal density throughout the depth of the cortex.

The profiles shown in Figure 4 and Supplementary Figure S1 also suggest that, in addition to the mean neuronal density, the variability of the neuronal density within an area may be another useful dimension to consider in order to understand patterns of change across the cortical surface. For example, the data for areas V3a and TEO showed considerable fluctuation around the mean, whereas the variations were much lower in area 6M. In order to capture this aspect of the data, we illustrate in Figure 8 the standard deviation of the neuronal density estimates within the counting frames assigned to each of the different areas. This analysis shows that visual, somatosensory, posterior parietal and caudal temporal areas tended to exhibit more variation in neuronal density through the cortical depth, when compared to motor, premotor, medial frontal and other temporal areas (note, for example, the low values obtained in subdivisions of the areas 4, 6, and 24 complexes, as well as in the agranular insula (AI), area 30, frontal areas 8b, 25 and 32, and caudal orbitofrontal areas OPAl and OPro). The overall neuronal density (Fig. 3) and the degree of cytoarchitectural differentiation in layers (Fig. 8) covaried significantly, but not in exact proportion: as shown in the insert in Figure 8 the coefficient of variation (ratio of the standard deviation to the mean) was far from uniform.

**Figure 8.**
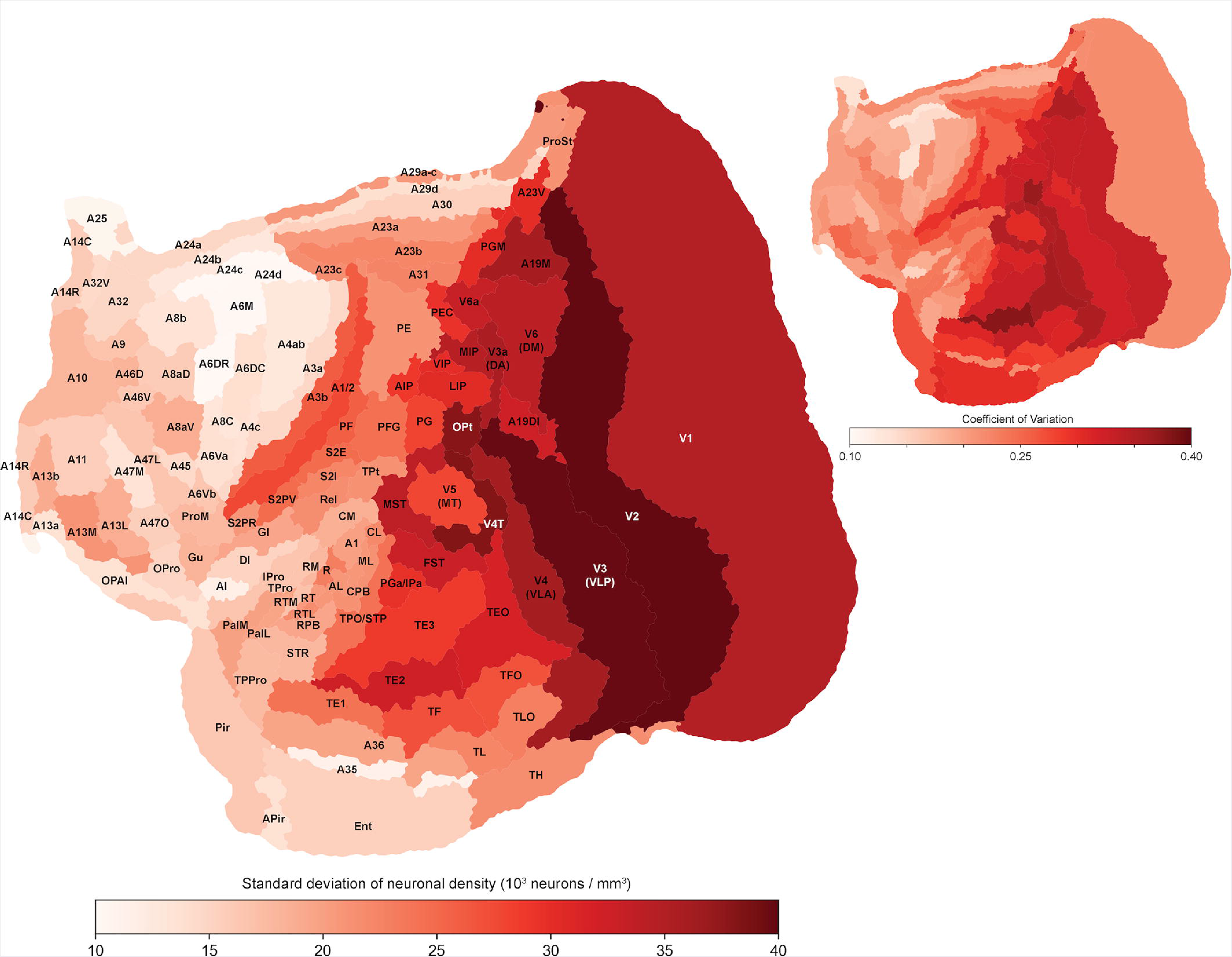
Standard deviation of neuronal density values obtained at multiple depths (in increments of 150 μm) in 4 locations of 116 areas, shown in an “unfolded” representation of the marmoset cortex (Majka et al. 2016). Areal boundaries were estimated using the criteria illustrated by Paxinos et al. (2012). Abbreviations are listed in Table 1. **Insert:** coefficient of variation of neuronal density for the same areas.

It has been suggested that regional variation in neuronal density across the cortex is primarily the result of changes in the upper (supragranular and granular) layers; in contrast, neuronal density in the infragranular layers would show a more modest degree of variation (Charvet et al. 2015). This conclusion was based on a grid of equally spaced locations along the anteroposterior and mediolateral axes. In order to test if this same effect was apparent in our area-based data, we calculated the ratio of neuronal density in the upper layers of the cortex (here approximated by the average density in the first two counting frames, which encompassed the top 300 μm of layers 2 and 3) versus lower layers (the first two complete counting frames above the limit of the white matter, which encompassed a similar volume of layers 5 and 6). This analysis, illustrated in Figure 9A, revealed significant relationships between the anteroposterior coordinate of an area and its neuronal density, both in the supragranular (r^2^= 0.486; F=107.8; p= 3.51·10^-18^) and infragranular (r^2^=0.071; F= 8.68; p= 3.89·10^-3^) layers. However, as suggested by Charvet et al. (2015), the variation was far more marked in the supragranular layers. The analysis illustrated in Figure 9B shows that the ratio of neuronal density in supragranular to infragranular layers varies significantly with anteroposterior location (r^2^=0.269; F=41.98; p=2.44·10^-9^).

**Figure 9:**
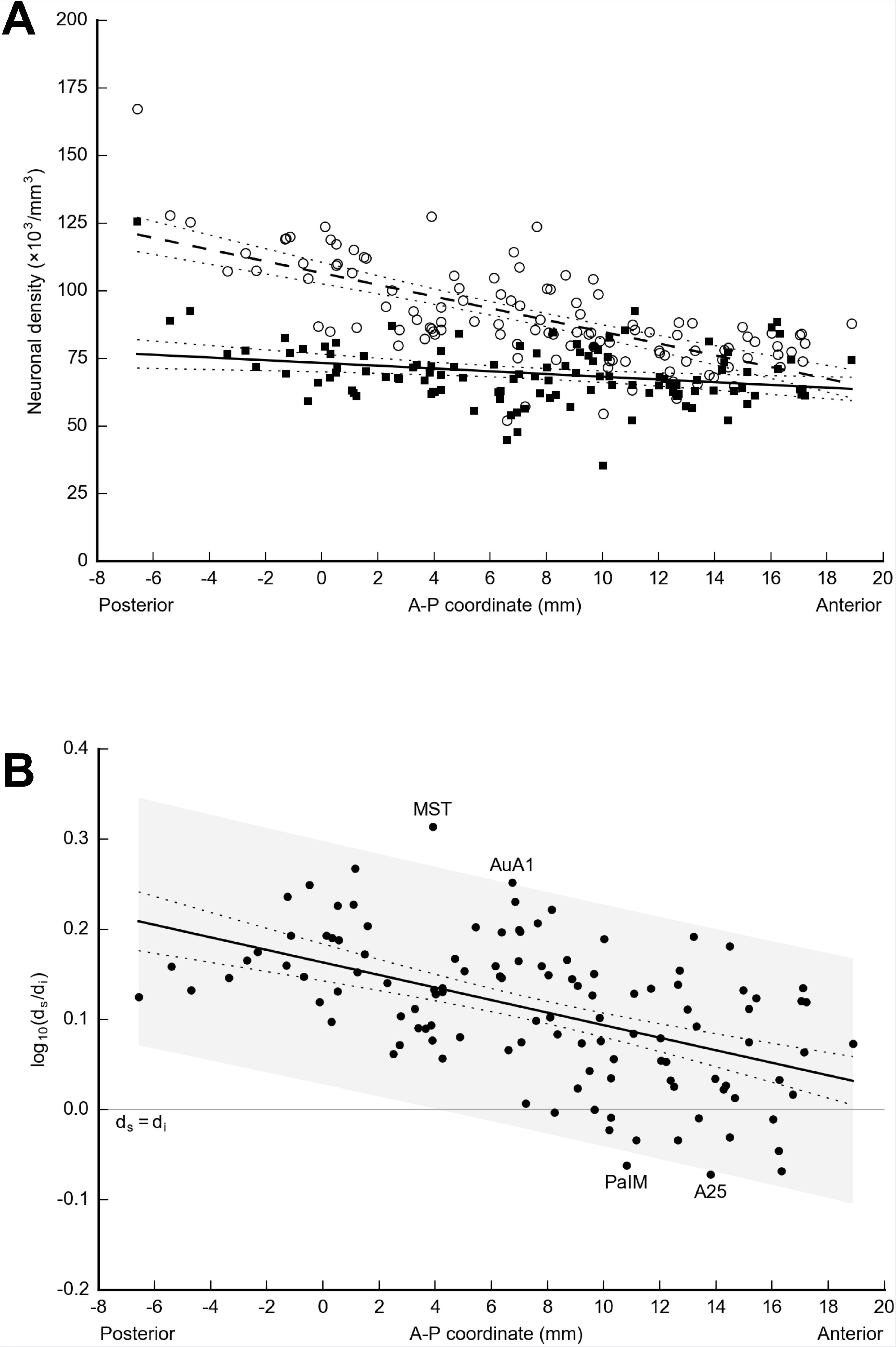
**A-** Neuronal density in the supragranular (white circles) and infragranular (black squares) layers, as a function of the A-P coordinate of the area’s barycenter. Linear regressions fitted to the data are shown in dashed and continuous lines, respectively, and the 95% confidence intervals for the regression slopes are shown by dotted lines. **B-** Logarithm of the ratio of neuronal densities in supragranular (d_s_) and infragranular (d_i_) layers of cortical areas, as a function of A-P coordinate. A linear model can account for most observations, with only 4 areas outside the 95% prediction interval.

### Cortical thickness and number of neurons per column

The thickness of the primate cortex is not uniform (e.g. Calabrese et al. 2015). Hence, two areas with identical mean neuronal density can have different numbers of neurons in a column extending from the pial surface to the white matter. These estimates are relevant in light of current models that attempt to correlate number of neurons in a column to the developmental gradient of neurogenesis along the dorsal and lateral surfaces of the primate brain (e.g. Calahane et al. 2012). In this section we report on calculations that provide analogous estimates for the marmoset cortex.

In order to correctly calculate the cortical thickness, we adopted a computational approach. The segmented cortex of animal CJ167 was reconstructed accurately in 3 dimensions, and radial “streamlines” were generated; the length of these streamlines accurately estimate the mean cortical thickness for the different areas (Figure 2; supplementary Figure S4). The results (Figure 10A) show that limbic areas near the corpus callosum (e.g. areas 24a, 25, 29a-c), ventral orbital surface (e.g. 13a, 13b, orbital periallocortex), and in the insular, parahippocampal, entorhinal and piriform regions tend to be thinner than isocortical areas on the lateral surface of the brain. It also supports the idea that mechanical factors related to the location of an area in highly convex or concave regions significantly affect cortical thickness (Hilgetag and Barbas 2006). For example, the lateral subdivision of area 47 (47L), the proisocortical motor area (ProM), subdivisions of the S2 complex, auditory core areas and the ventral subdivision area 23 (23V) tend to be thicker than adjacent areas, whereas polysensory area PGa/IPa, which follows he fundus of the superior temporal sulcus, is markedly thinner than adjacent areas. Perhaps more surprisingly, this analysis also shows that areas of the posterior parietal cortex (e.g. AIP, LIP, VIP and inferior parietal areas such as PG and PFG) are markedly thicker than adjacent visual and somatosensory areas, despite being located in relatively “flat” regions of cortex. Although our data show some evidence of the trend detected by Calahane et al. (2012), whereby the isocortex in the lateral surface of the cortex (excluding limbic regions) tends to become thicker along a gradient running approximately ventrocaudal to rostral, this relationship is not strong (Figure 10 and Supplementary Table S2).

**Figure 10.**
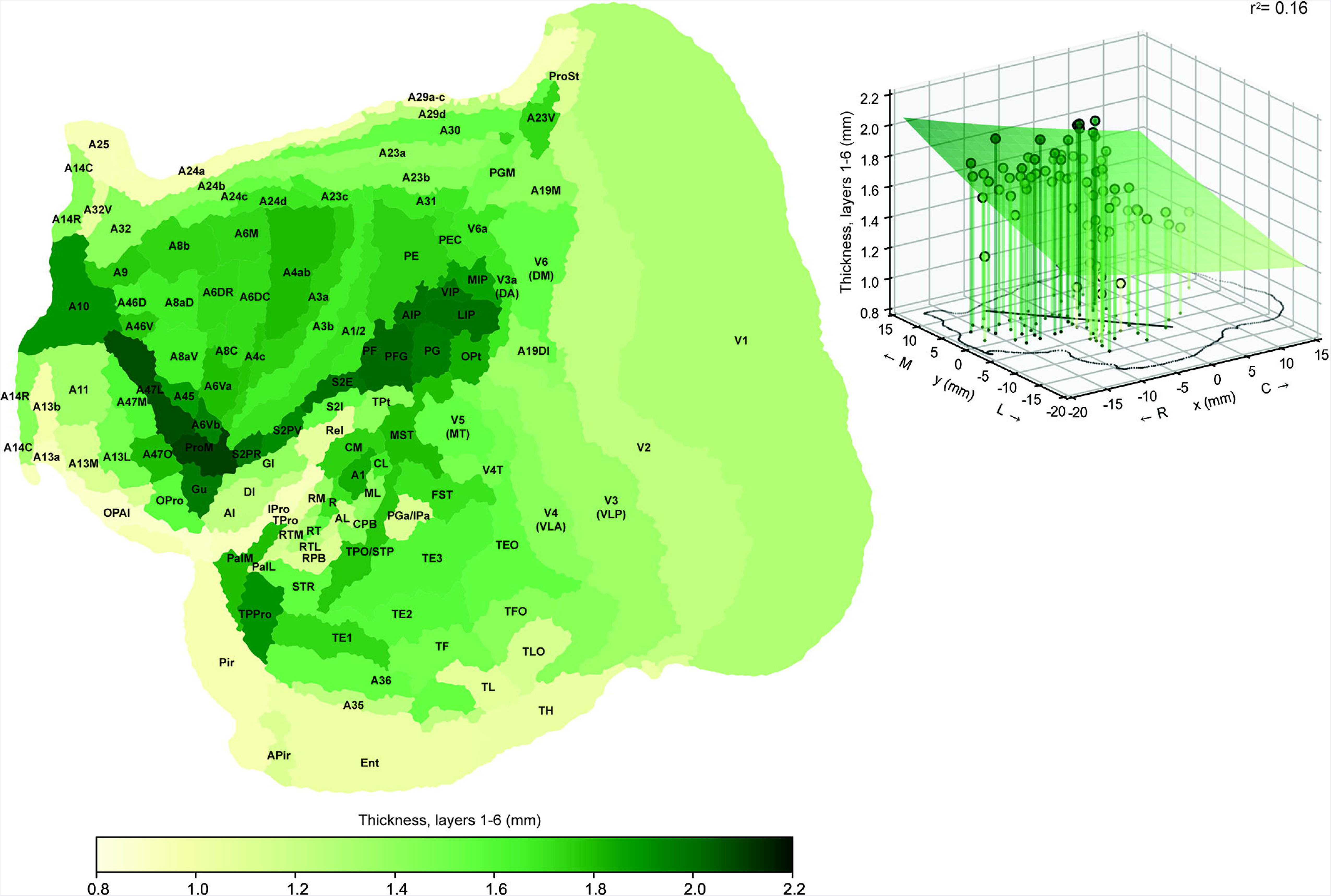
Mean cortical thickness for 116 cytoarchitectural areas of the marmoset brain, shown in an “unfolded” representation of the cortex (Majka et al. 2016). Areal boundaries were estimated using the criteria illustrated by Paxinos et al. (2012). Abbreviations are listed in Table 1. **Insert:** 3-dimensional plot of cortical thickness according to mediolateral (M-L) and rostrocaudal (R-C) location of the barycentre of each cortical area. The outline of a flat map of the marmoset cortex is shown in the horizontal plane (see Figure 3 for a similar map showing cortical areas). In this analysis, only isocortical areas were used, similar to Calahane et al. (2012) and Charvet et al. (2015). As in these studies, to identify the primary direction of variation (“primary axis”), we fitted a 2-dimensional surface to the cartesian coordinates of the barycentres and the values of the cortical thickness of individual areas. The principal axis is indicated by the thick line overlying the flat map. The list of areas used in this analysis is given in Supplementary Table S1, and the parameters of the fitted surface are provided in Supplementary Table S2.

Based on the thickness of the cortex, and measurements of mean neuronal density for layer 1 and for layers 2-6 (Tables 1 and 2), we calculated the mean number of neurons under a columnar compartment with a cross-section of 1 mm^2^, measured parallel to the cortical surface (for convenience, we will refer to such compartments as “unit columns”). The results for different areas, illustrated in Figure 11 and detailed in Table 3, show that variations in the number of neurons per unit column resemble those of neuronal density in some, but not all aspects. For example, as for neuronal density, the primary visual and somatosensory areas (V1 and area 3b) show higher estimates of numbers of neurons per unit column in comparison with adjacent areas. However, in the auditory cortex one finds a distinct pattern, whereby the core (A1, R, RT) and caudal belt areas (CL, CM) show the highest numbers of neurons per unit column, and the RPB the lowest. Thus, variations in cortical thickness can offset gradients of neuronal density, resulting in a distinctive spatial pattern. In visual cortex, the ordered caudal to rostral trends revealed by the analysis of neuronal density become less obvious. Instead, one finds similar estimates of numbers of neurons per unit column in areas V2, V4, TEO, and TE1-3, while visuomotor areas in the posterior parietal (AIP, LIP, MIP, OPt, VIP and V6a) and superior temporal (MST, FST) regions stand out by virtue of high numbers of neurons per unit column. Some of the other extrastriate areas that contribute to the “dorsal stream” also appear to have moderately increased numbers of cells per column in comparison with ventral stream areas (e.g. V3a, V4t, V6). In addition, other higher-order association areas in the precuneus (23V, PGM), superior temporal cortex (TPO/STP), temporal pole (TPPro) and ventrolateral prefrontal cortex (areas 10 and 47L) show relatively high numbers of neurons in a column, in comparison with most sensory and motor areas. Fitting a model to the data obtained for isocortical areas (Fig. 11) reveals a significant relationship, which, as in other primates (Cahalane et al. 2012), follows a principal axis running from posterior and medial areas to anterior and lateral areas. However, the correlation (r^2^= 0.28; Supplementary Table S2) is much lower than that observed in the analysis of neuronal density using the same set of areas (r^2^= 0.78; Supplementary Figure S5).

**Figure 11.**
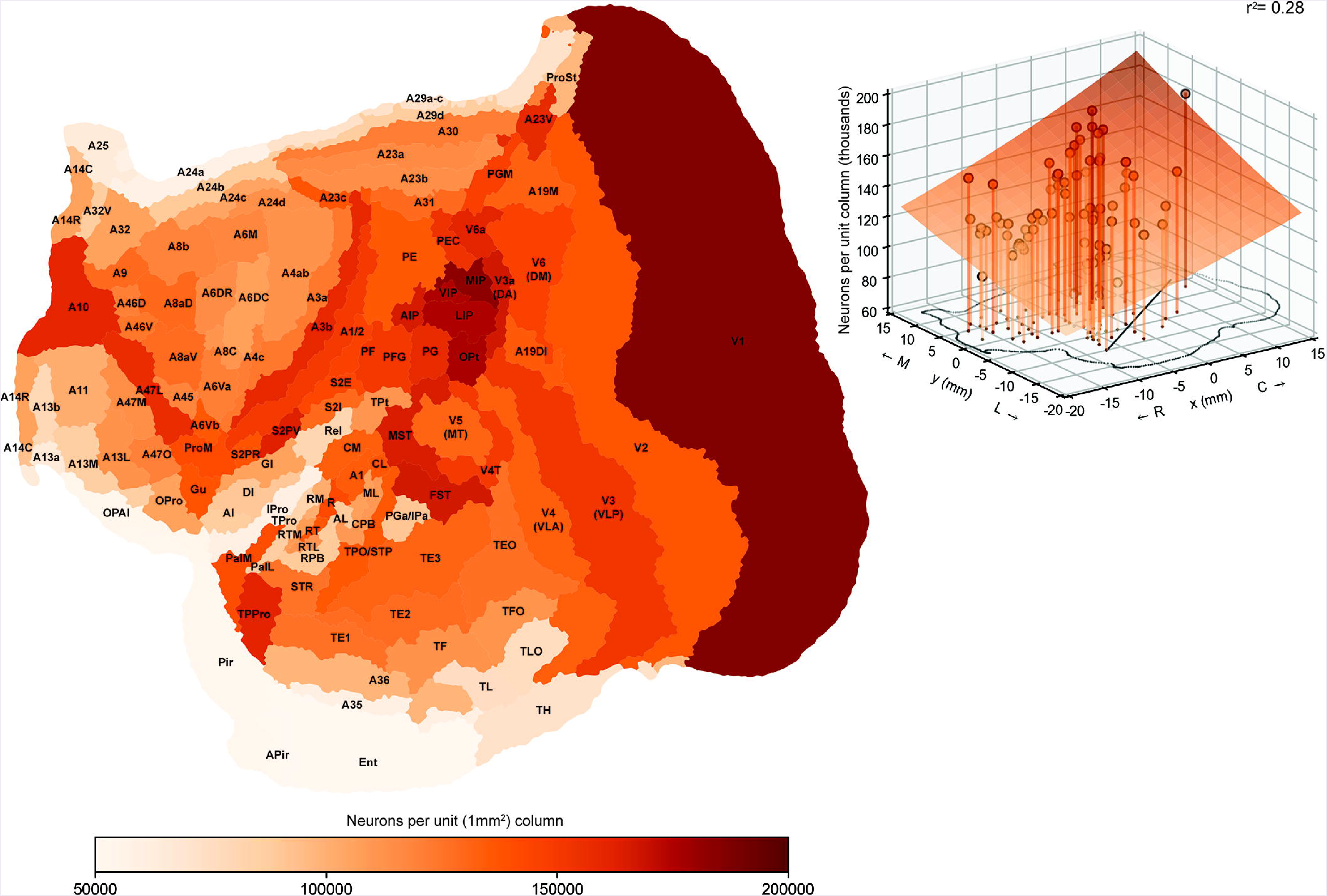
Number of neurons under a columnar compartment of cortex with a 1 mm^2^ cross-section (unit column), shown in an “unfolded” representation of the cortex (Majka et al. 2016). Areal boundaries were estimated using the criteria illustrated by Paxinos et al. (2012). Abbreviations are listed in Table 1. **Insert:** 3-dimensional plot according to mediolateral (M-L) and rostrocaudal (R-C) location of the barycentre of each cortical area. The outline of a flat map of the marmoset cortex is shown in the horizontal plane. For details, see legend of Figure 10.

**Table 3:**
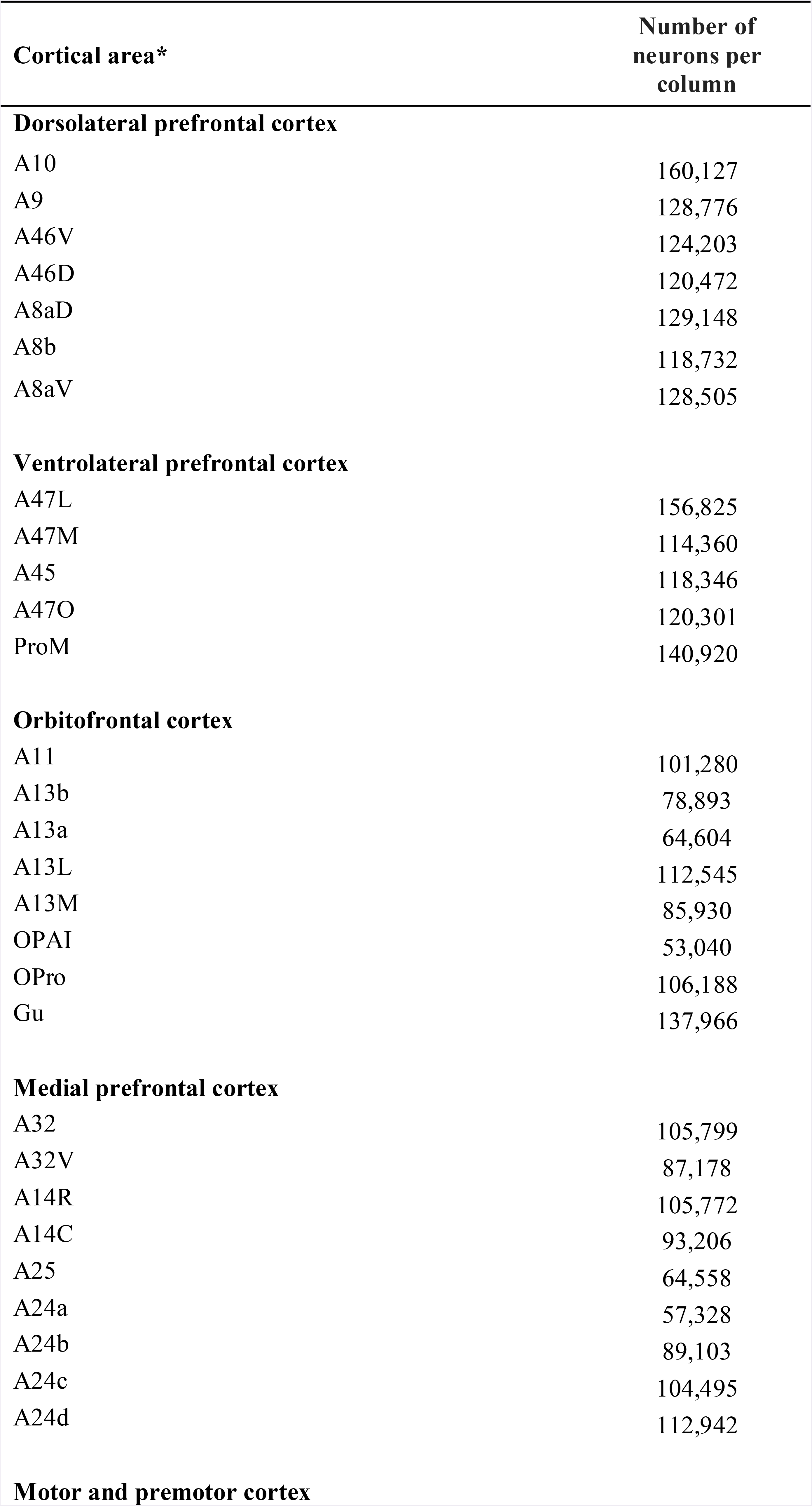
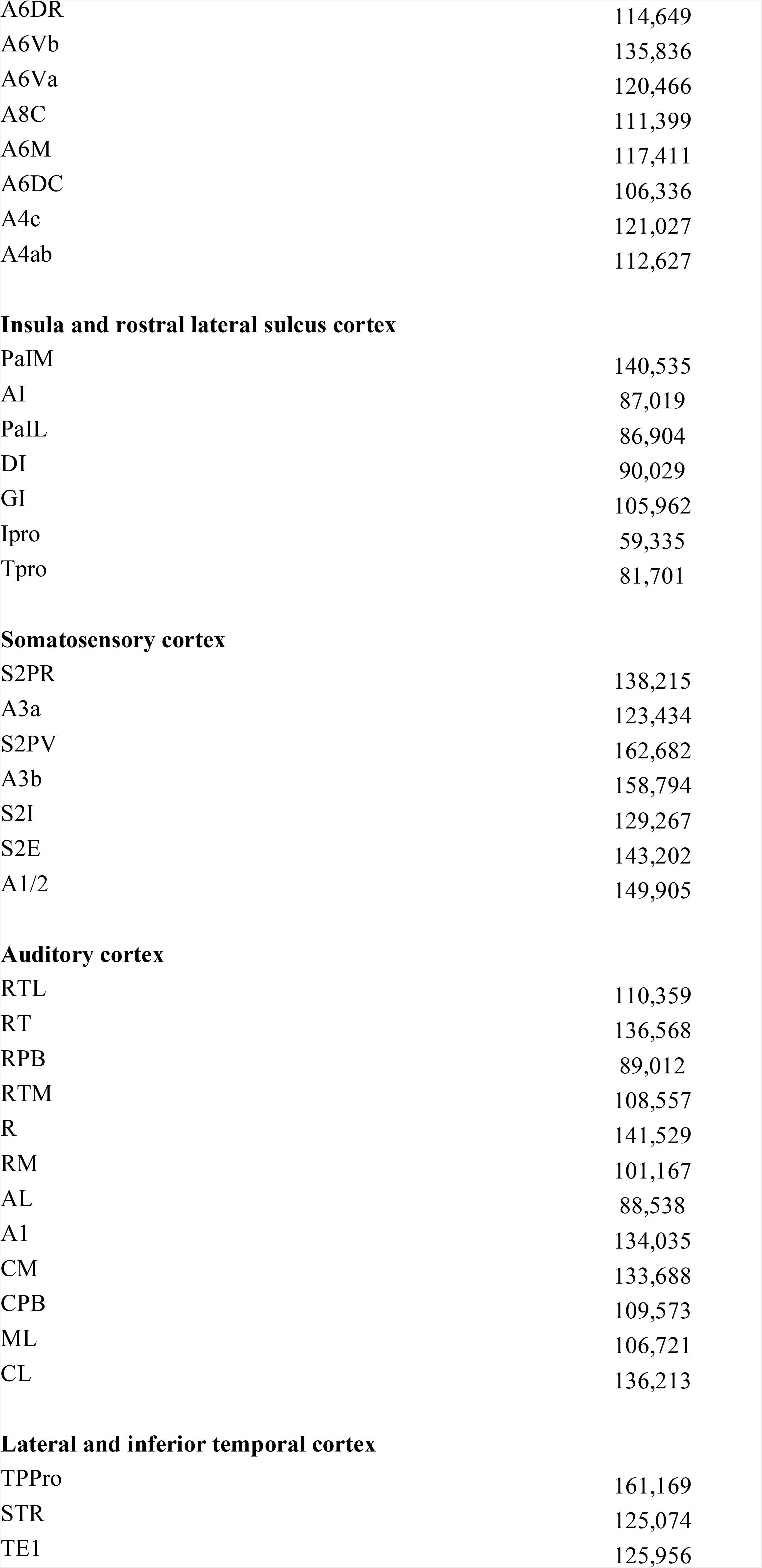
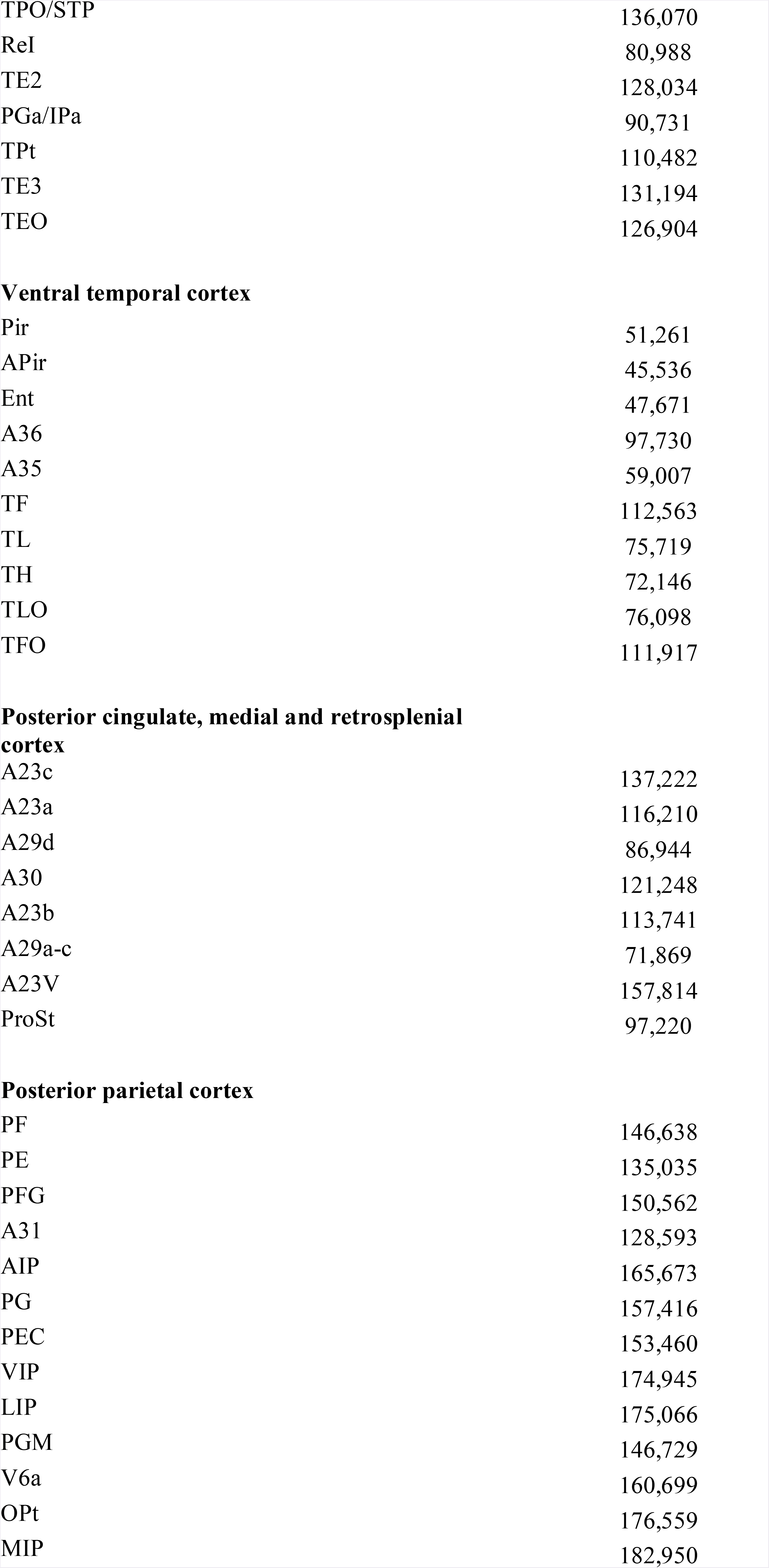
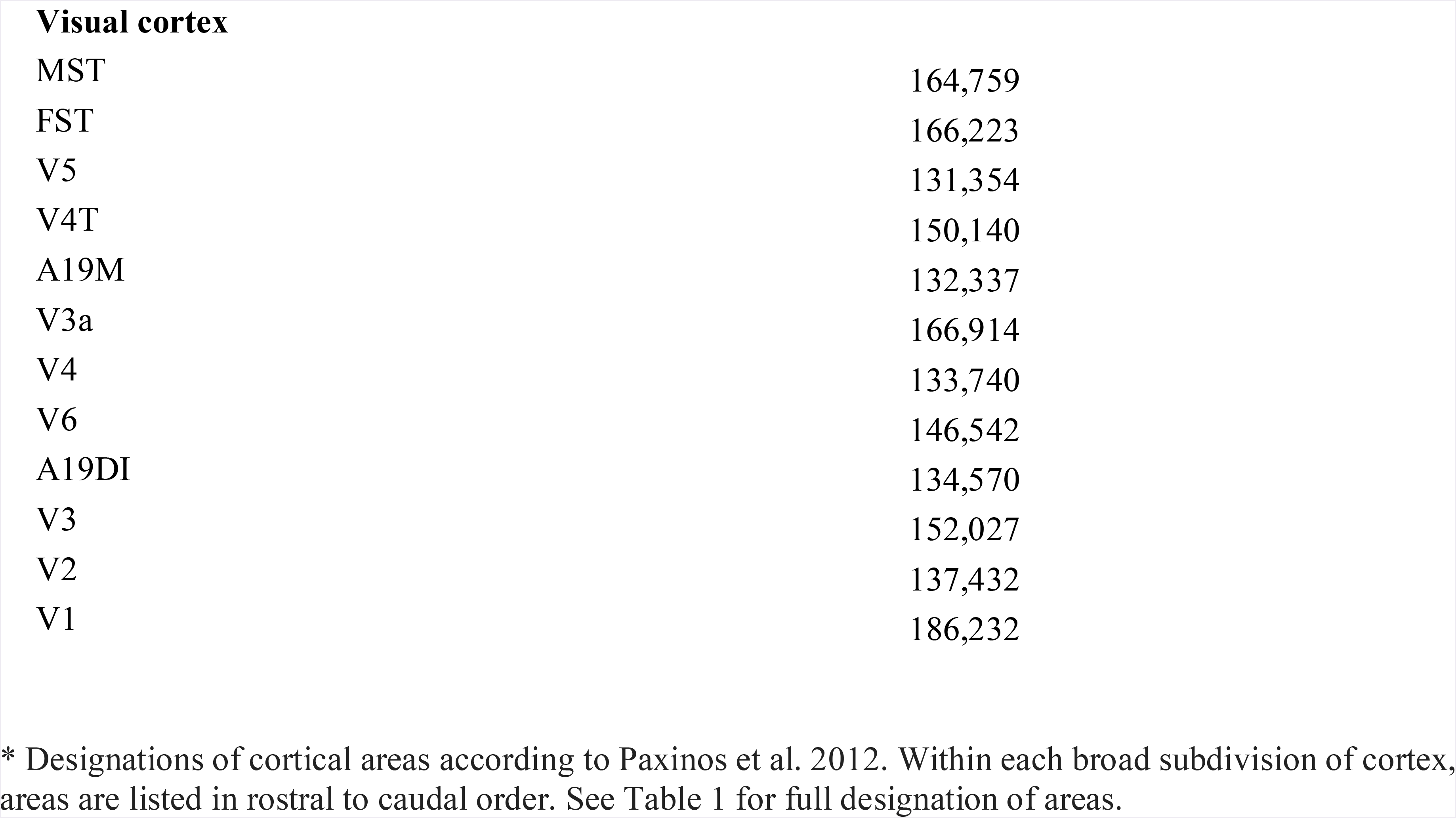
Number of neurons within a column with a cross-section of 1mm^2^ parallel to the cortical surface

| **Cortical area\*** | **Number of neurons per column** |
| --- | --- |
| **Dorsolateral prefrontal cortex** |  |
| A10 | 160,127 |
| A9 | 128,776 |
| A46V | 124,203 |
| A46D | 120,472 |
| A8aD | 129,148 |
| A8b | 118,732 |
| A8aV | 128,505 |
| **Ventrolateral prefrontal cortex** |  |
| A47L | 156,825 |
| A47M | 114,360 |
| A45 | 118,346 |
| A47O | 120,301 |
| ProM | 140,920 |
| **Orbitofrontal cortex** |  |
| A11 | 101,280 |
| A13b | 78,893 |
| A13a | 64,604 |
| A13L | 112,545 |
| A13M | 85,930 |
| OPAI | 53,040 |
| OPro | 106,188 |
| Gu | 137,966 |
| **Medial prefrontal cortex** |  |
| A32 | 105,799 |
| A32V | 87,178 |
| A14R | 105,772 |
| A14C | 93,206 |
| A25 | 64,558 |
| A24a | 57,328 |
| A24b | 89,103 |
| A24c | 104,495 |
| A24d | 112,942 |
| **Motor and premotor cortex** |  |
| A6DR | 114,649 |
| A6Vb | 135,836 |
| A6Va | 120,466 |
| A8C | 111,399 |
| A6M | 117,411 |
| A6DC | 106,336 |
| A4c | 121,027 |
| A4ab | 112,627 |
| **Insula and rostral lateral sulcus cortex** |  |
| PaIM | 140,535 |
| AI | 87,019 |
| PaIL | 86,904 |
| DI | 90,029 |
| GI | 105,962 |
| Ipro | 59,335 |
| Tpro | 81,701 |
| **Somatosensory cortex** |  |
| S2PR | 138,215 |
| A3a | 123,434 |
| S2PV | 162,682 |
| A3b | 158,794 |
| S2I | 129,267 |
| S2E | 143,202 |
| A1/2 | 149,905 |
| **Auditory cortex** |  |
| RTL | 110,359 |
| RT | 136,568 |
| RPB | 89,012 |
| RTM | 108,557 |
| R | 141,529 |
| RM | 101,167 |
| AL | 88,538 |
| A1 | 134,035 |
| CM | 133,688 |
| CPB | 109,573 |
| ML | 106,721 |
| CL | 136,213 |
| **Lateral and inferior temporal cortex** |  |
| TPPro | 161,169 |
| STR | 125,074 |
| TE1 | 125,956 |
| TPO/STP | 136,070 |
| ReI | 80,988 |
| TE2 | 128,034 |
| PGa/IPa | 90,731 |
| TPt | 110,482 |
| TE3 | 131,194 |
| TEO | 126,904 |
| **Ventral temporal cortex** |  |
| Pir | 51,261 |
| APir | 45,536 |
| Ent | 47,671 |
| A36 | 97,730 |
| A35 | 59,007 |
| TF | 112,563 |
| TL | 75,719 |
| TH | 72,146 |
| TLO | 76,098 |
| TFO | 111,917 |
| **Posterior cingulate, medial and retrosplenial cortex** |  |
| A23c | 137,222 |
| A23a | 116,210 |
| A29d | 86,944 |
| A30 | 121,248 |
| A23b | 113,741 |
| A29a-c | 71,869 |
| A23V | 157,814 |
| ProSt | 97,220 |
| **Posterior parietal cortex** |  |
| PF | 146,638 |
| PE | 135,035 |
| PFG | 150,562 |
| A31 | 128,593 |
| AIP | 165,673 |
| PG | 157,416 |
| PEC | 153,460 |
| VIP | 174,945 |
| LIP | 175,066 |
| PGM | 146,729 |
| V6a | 160,699 |
| OPt | 176,559 |
| MIP | 182,950 |
| **Visual cortex** |  |
| MST | 164,759 |
| FST | 166,223 |
| V5 | 131,354 |
| V4T | 150,140 |
| A19M | 132,337 |
| V3a | 166,914 |
| V4 | 133,740 |
| V6 | 146,542 |
| A19DI | 134,570 |
| V3 | 152,027 |
| V2 | 137,432 |
| V1 | 186,232 |

\* Designations of cortical areas according to Paxinos et al. 2012. Within each broad subdivision of cortex, areas are listed in rostral to caudal order. See Table 1 for full designation of areas.

### Volumes of areas and neuronal counts

Finally, the accurate computational reconstruction of the cortex, combined with the analysis of neuronal density, allowed us to estimate the volumes and numbers of neurons within different cortical areas of the marmoset (Table 4). Overall, the cortex of marmoset CJ167 (excluding the hippocampal formation) was estimated to occupy a volume of 1584 mm^3^ (corrected for shrinkage), and to contain approximately 1.5 × 10^8^ neurons in each hemisphere. This analysis revealed that area V1 alone occupies 18.8% of the volume of the cortex, and contains a staggering 28.6% of the cortical neurons. Moreover, the classical striate and extrastriate areas (grouped as “visual cortex”, in Table 4) together account for 40.8% of the volume of the cortex, and contain 52.8% of the neurons. In comparison, auditory areas (core, belt and parabelt) contain 2.5% of the cortical neurons, and somatosensory areas (encompassing all subdivisions of the classical “S1” and “S2” complexes) another 4.9%. Groupings such as those used in the present tables reflect a somewhat artificial attempt to classify areas according to location and function; for example, areas TE and TEO could arguably be appropriately classified as “visual cortex”, the superior temporal rostral cortex (STR) as “auditory”, and posterior parietal areas PE and PF as “somatosensory”. Nonetheless, the analysis presented in Table 4, highlights the heavy dominance of vision as the primary sensory modality in primates.

**Table 4:**
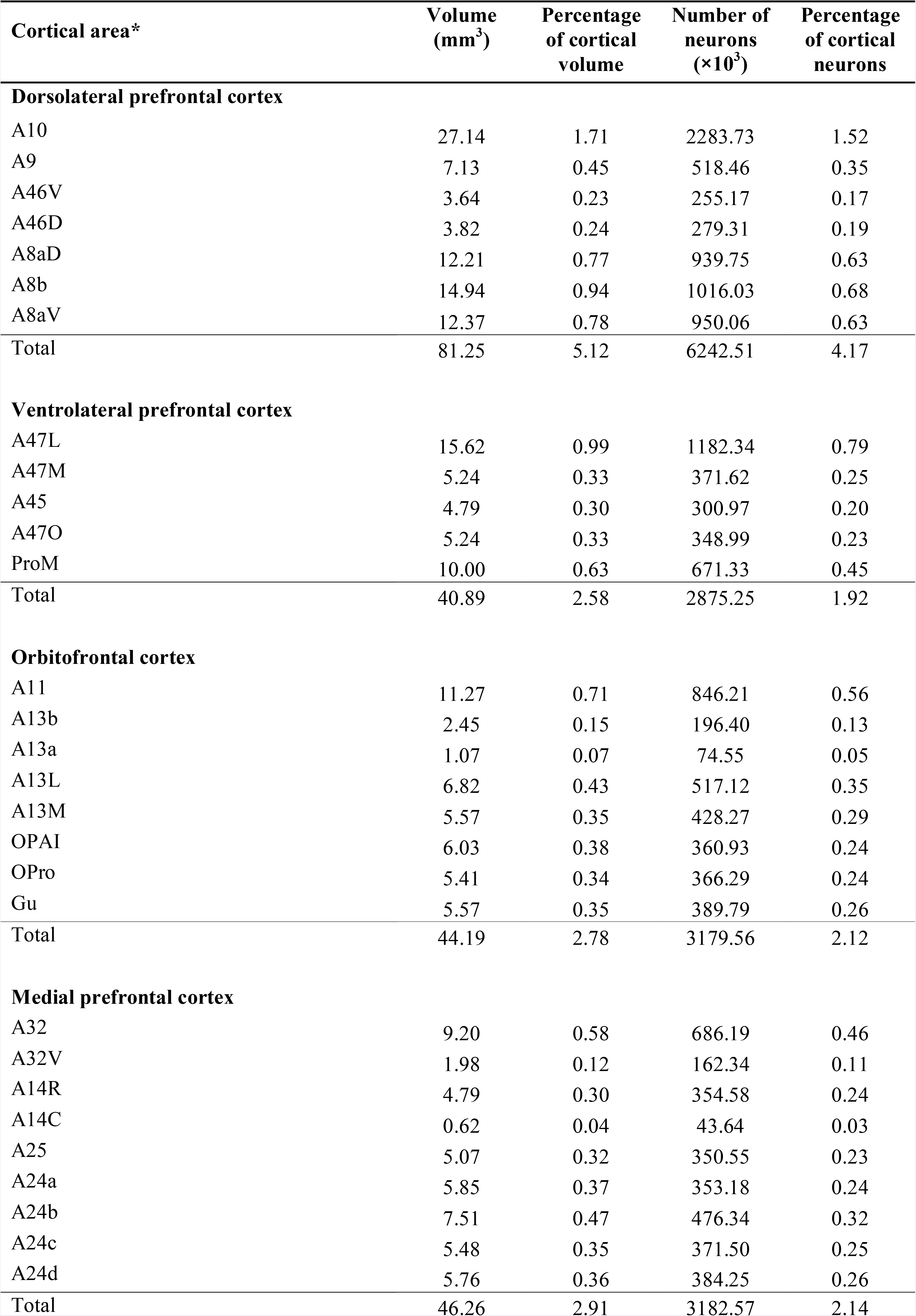
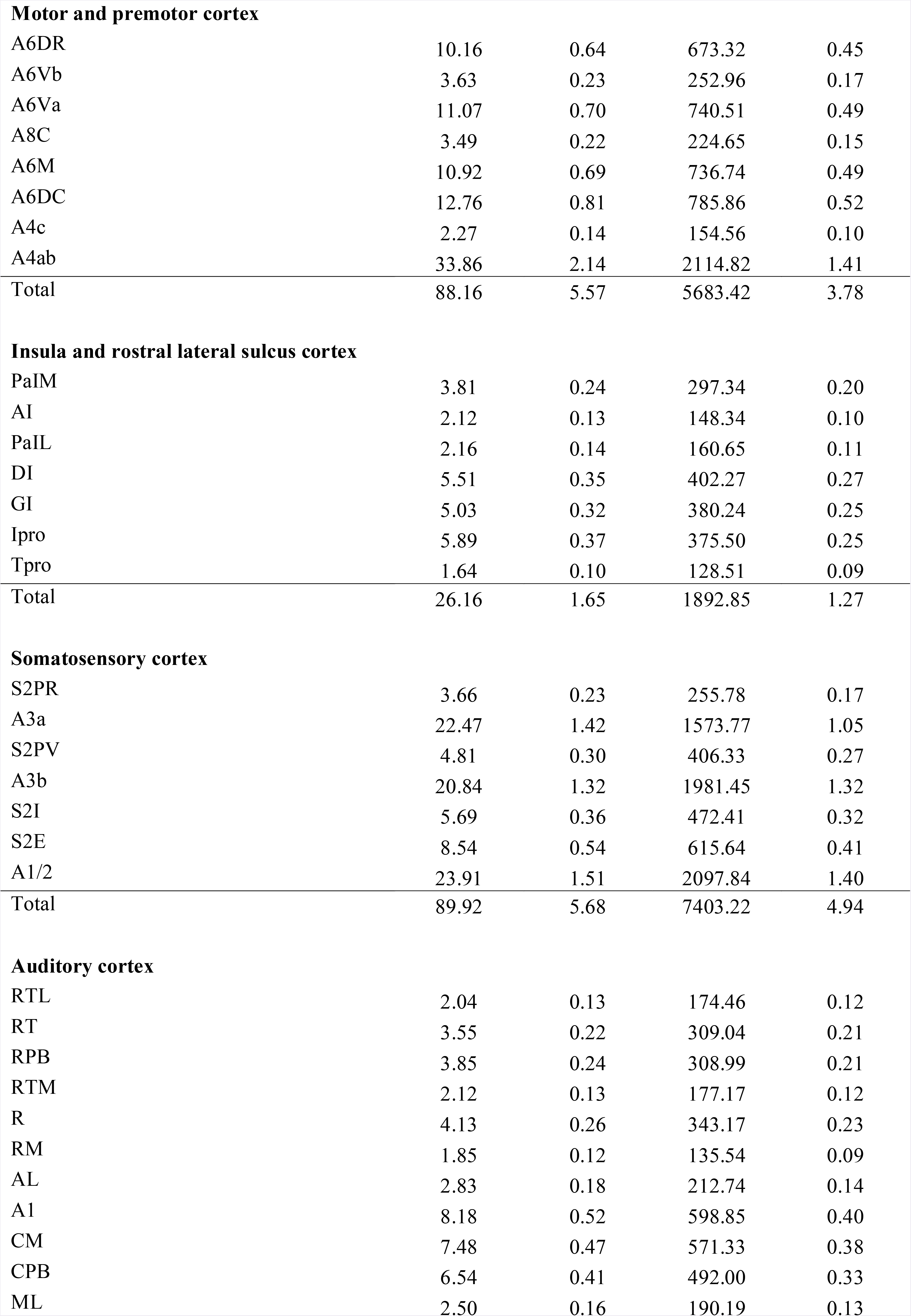
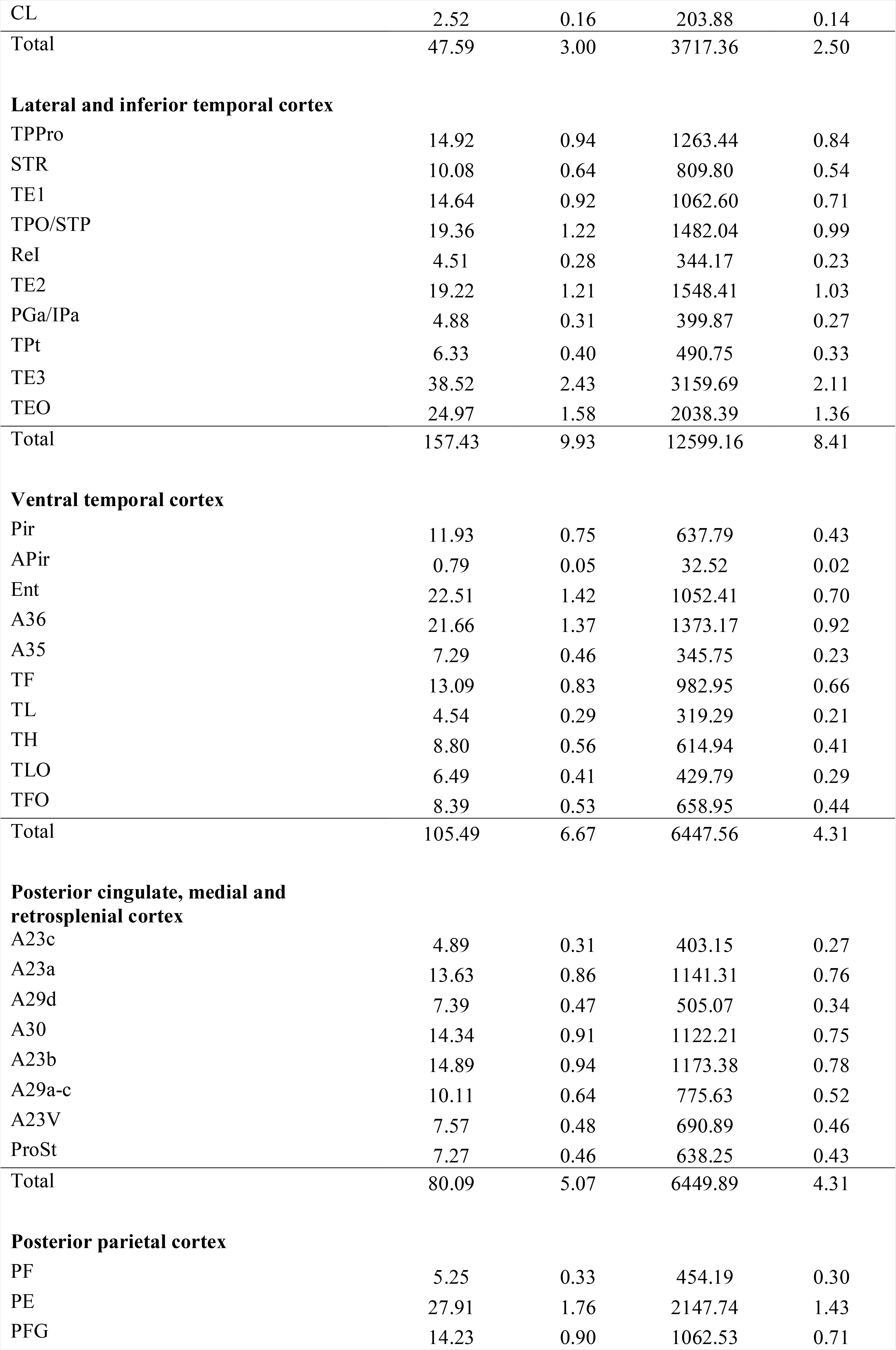
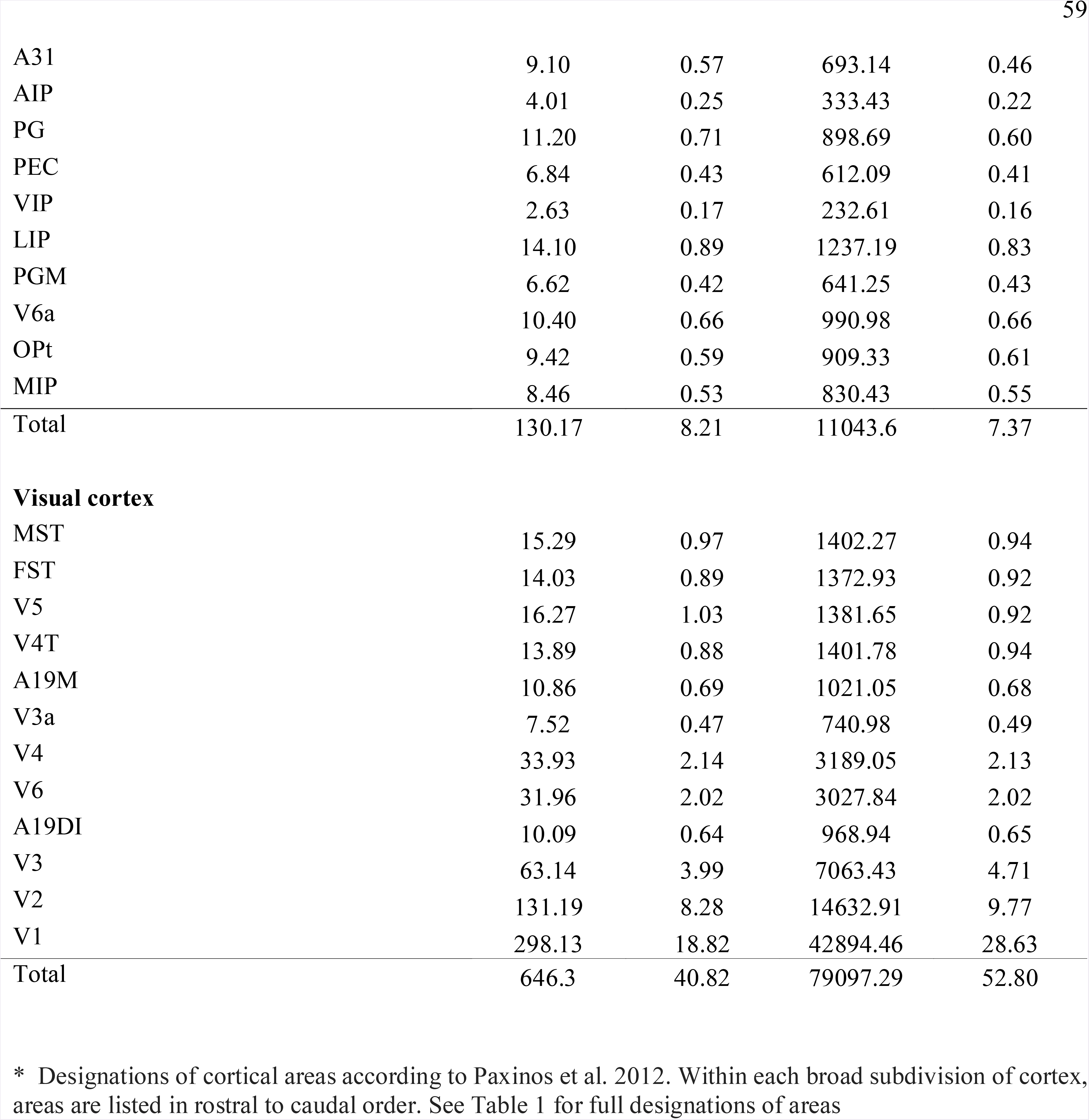
Volumes of areas and number of neurons in one hemisphere of the marmoset cortex

| **Cortical area\*** | **Volume**<br>**(mm<sup>3</sup>)** | **Percentage**<br>**of cortical**<br>**volume** | **Number of**<br>**neurons**<br>**(×10<sup>3</sup>)** | **Percentage**<br>**of cortical**<br>**neurons** |
| --- | --- | --- | --- | --- |
| **Dorsolateral prefrontal cortex** |  |  |  |  |
| A10 | 27.14 | 1.71 | 2283.73 | 1.52 |
| A9 | 7.13 | 0.45 | 518.46 | 0.35 |
| A46V | 3.64 | 0.23 | 255.17 | 0.17 |
| A46D | 3.82 | 0.24 | 279.31 | 0.19 |
| A8aD | 12.21 | 0.77 | 939.75 | 0.63 |
| A8b | 14.94 | 0.94 | 1016.03 | 0.68 |
| A8aV | 12.37 | 0.78 | 950.06 | 0.63 |
| Total | 81.25 | 5.12 | 6242.51 | 4.17 |
| **Ventrolateral prefrontal cortex** |  |  |  |  |
| A47L | 15.62 | 0.99 | 1182.34 | 0.79 |
| A47M | 5.24 | 0.33 | 371.62 | 0.25 |
| A45 | 4.79 | 0.30 | 300.97 | 0.20 |
| A47O | 5.24 | 0.33 | 348.99 | 0.23 |
| ProM | 10.00 | 0.63 | 671.33 | 0.45 |
| Total | 40.89 | 2.58 | 2875.25 | 1.92 |
| **Orbitofrontal cortex** |  |  |  |  |
| A11 | 11.27 | 0.71 | 846.21 | 0.56 |
| A13b | 2.45 | 0.15 | 196.40 | 0.13 |
| A13a | 1.07 | 0.07 | 74.55 | 0.05 |
| A13L | 6.82 | 0.43 | 517.12 | 0.35 |
| A13M | 5.57 | 0.35 | 428.27 | 0.29 |
| OPAI | 6.03 | 0.38 | 360.93 | 0.24 |
| OPro | 5.41 | 0.34 | 366.29 | 0.24 |
| Gu | 5.57 | 0.35 | 389.79 | 0.26 |
| Total | 44.19 | 2.78 | 3179.56 | 2.12 |
| **Medial prefrontal cortex** |  |  |  |  |
| A32 | 9.20 | 0.58 | 686.19 | 0.46 |
| A32V | 1.98 | 0.12 | 162.34 | 0.11 |
| A14R | 4.79 | 0.30 | 354.58 | 0.24 |
| A14C | 0.62 | 0.04 | 43.64 | 0.03 |
| A25 | 5.07 | 0.32 | 350.55 | 0.23 |
| A24a | 5.85 | 0.37 | 353.18 | 0.24 |
| A24b | 7.51 | 0.47 | 476.34 | 0.32 |
| A24c | 5.48 | 0.35 | 371.50 | 0.25 |
| A24d | 5.76 | 0.36 | 384.25 | 0.26 |
| Total | 46.26 | 2.91 | 3182.57 | 2.14 |
| **Motor and premotor cortex** |  |  |  |  |
| A6DR | 10.16 | 0.64 | 673.32 | 0.45 |
| A6Vb | 3.63 | 0.23 | 252.96 | 0.17 |
| A6Va | 11.07 | 0.70 | 740.51 | 0.49 |
| A8C | 3.49 | 0.22 | 224.65 | 0.15 |
| A6M | 10.92 | 0.69 | 736.74 | 0.49 |
| A6DC | 12.76 | 0.81 | 785.86 | 0.52 |
| A4c | 2.27 | 0.14 | 154.56 | 0.10 |
| A4ab | 33.86 | 2.14 | 2114.82 | 1.41 |
| Total | 88.16 | 5.57 | 5683.42 | 3.78 |
| **Insula and rostral lateral sulcus cortex** |  |  |  |  |
| PaIM | 3.81 | 0.24 | 297.34 | 0.20 |
| AI | 2.12 | 0.13 | 148.34 | 0.10 |
| PaIL | 2.16 | 0.14 | 160.65 | 0.11 |
| DI | 5.51 | 0.35 | 402.27 | 0.27 |
| GI | 5.03 | 0.32 | 380.24 | 0.25 |
| Ipro | 5.89 | 0.37 | 375.50 | 0.25 |
| Tpro | 1.64 | 0.10 | 128.51 | 0.09 |
| Total | 26.16 | 1.65 | 1892.85 | 1.27 |
| **Somatosensory cortex** |  |  |  |  |
| S2PR | 3.66 | 0.23 | 255.78 | 0.17 |
| A3a | 22.47 | 1.42 | 1573.77 | 1.05 |
| S2PV | 4.81 | 0.30 | 406.33 | 0.27 |
| A3b | 20.84 | 1.32 | 1981.45 | 1.32 |
| S2I | 5.69 | 0.36 | 472.41 | 0.32 |
| S2E | 8.54 | 0.54 | 615.64 | 0.41 |
| A1/2 | 23.91 | 1.51 | 2097.84 | 1.40 |
| Total | 89.92 | 5.68 | 7403.22 | 4.94 |
| **Auditory cortex** |  |  |  |  |
| RTL | 2.04 | 0.13 | 174.46 | 0.12 |
| RT | 3.55 | 0.22 | 309.04 | 0.21 |
| RPB | 3.85 | 0.24 | 308.99 | 0.21 |
| RTM | 2.12 | 0.13 | 177.17 | 0.12 |
| R | 4.13 | 0.26 | 343.17 | 0.23 |
| RM | 1.85 | 0.12 | 135.54 | 0.09 |
| AL | 2.83 | 0.18 | 212.74 | 0.14 |
| A1 | 8.18 | 0.52 | 598.85 | 0.40 |
| CM | 7.48 | 0.47 | 571.33 | 0.38 |
| CPB | 6.54 | 0.41 | 492.00 | 0.33 |
| ML | 2.50 | 0.16 | 190.19 | 0.13 |
| CL | 2.52 | 0.16 | 203.88 | 0.14 |
| Total | 47.59 | 3.00 | 3717.36 | 2.50 |
| **Lateral and inferior temporal cortex** |  |  |  |  |
| TPPro | 14.92 | 0.94 | 1263.44 | 0.84 |
| STR | 10.08 | 0.64 | 809.80 | 0.54 |
| TE1 | 14.64 | 0.92 | 1062.60 | 0.71 |
| TPO/STP | 19.36 | 1.22 | 1482.04 | 0.99 |
| ReI | 4.51 | 0.28 | 344.17 | 0.23 |
| TE2 | 19.22 | 1.21 | 1548.41 | 1.03 |
| PGa/IPa | 4.88 | 0.31 | 399.87 | 0.27 |
| TPt | 6.33 | 0.40 | 490.75 | 0.33 |
| TE3 | 38.52 | 2.43 | 3159.69 | 2.11 |
| TEO | 24.97 | 1.58 | 2038.39 | 1.36 |
| Total | 157.43 | 9.93 | 12599.16 | 8.41 |
| **Ventral temporal cortex** |  |  |  |  |
| Pir | 11.93 | 0.75 | 637.79 | 0.43 |
| APir | 0.79 | 0.05 | 32.52 | 0.02 |
| Ent | 22.51 | 1.42 | 1052.41 | 0.70 |
| A36 | 21.66 | 1.37 | 1373.17 | 0.92 |
| A35 | 7.29 | 0.46 | 345.75 | 0.23 |
| TF | 13.09 | 0.83 | 982.95 | 0.66 |
| TL | 4.54 | 0.29 | 319.29 | 0.21 |
| TH | 8.80 | 0.56 | 614.94 | 0.41 |
| TLO | 6.49 | 0.41 | 429.79 | 0.29 |
| TFO | 8.39 | 0.53 | 658.95 | 0.44 |
| Total | 105.49 | 6.67 | 6447.56 | 4.31 |
| **Posterior cingulate, medial and retrosplenial cortex** |  |  |  |  |
| A23c | 4.89 | 0.31 | 403.15 | 0.27 |
| A23a | 13.63 | 0.86 | 1141.31 | 0.76 |
| A29d | 7.39 | 0.47 | 505.07 | 0.34 |
| A30 | 14.34 | 0.91 | 1122.21 | 0.75 |
| A23b | 14.89 | 0.94 | 1173.38 | 0.78 |
| A29a-c | 10.11 | 0.64 | 775.63 | 0.52 |
| A23V | 7.57 | 0.48 | 690.89 | 0.46 |
| ProSt | 7.27 | 0.46 | 638.25 | 0.43 |
| Total | 80.09 | 5.07 | 6449.89 | 4.31 |
| **Posterior parietal cortex** |  |  |  |  |
| PF | 5.25 | 0.33 | 454.19 | 0.30 |
| PE | 27.91 | 1.76 | 2147.74 | 1.43 |
| PFG | 14.23 | 0.90 | 1062.53 | 0.71 |
| A31 | 9.10 | 0.57 | 693.14 | 0.46 |
| AIP | 4.01 | 0.25 | 333.43 | 0.22 |
| PG | 11.20 | 0.71 | 898.69 | 0.60 |
| PEC | 6.84 | 0.43 | 612.09 | 0.41 |
| VIP | 2.63 | 0.17 | 232.61 | 0.16 |
| LIP | 14.10 | 0.89 | 1237.19 | 0.83 |
| PGM | 6.62 | 0.42 | 641.25 | 0.43 |
| V6a | 10.40 | 0.66 | 990.98 | 0.66 |
| OPt | 9.42 | 0.59 | 909.33 | 0.61 |
| MIP | 8.46 | 0.53 | 830.43 | 0.55 |
| Total | 130.17 | 8.21 | 11043.6 | 7.37 |
| **Visual cortex** |  |  |  |  |
| MST | 15.29 | 0.97 | 1402.27 | 0.94 |
| FST | 14.03 | 0.89 | 1372.93 | 0.92 |
| V5 | 16.27 | 1.03 | 1381.65 | 0.92 |
| V4T | 13.89 | 0.88 | 1401.78 | 0.94 |
| A19M | 10.86 | 0.69 | 1021.05 | 0.68 |
| V3a | 7.52 | 0.47 | 740.98 | 0.49 |
| V4 | 33.93 | 2.14 | 3189.05 | 2.13 |
| V6 | 31.96 | 2.02 | 3027.84 | 2.02 |
| A19DI | 10.09 | 0.64 | 968.94 | 0.65 |
| V3 | 63.14 | 3.99 | 7063.43 | 4.71 |
| V2 | 131.19 | 8.28 | 14632.91 | 9.77 |
| V1 | 298.13 | 18.82 | 42894.46 | 28.63 |
| Total | 646.3 | 40.82 | 79097.29 | 52.80 |

\* Designations of cortical areas according to Paxinos et al. 2012. Within each broad subdivision of cortex, areas are listed in rostral to caudal order. See Table 1 for full designations of areas

## Discussion

We report on neuronal density values for each of the currently recognized cortical areas of the marmoset monkey, obtained through the application of stereological methods to sections stained for a specific neuronal marker (NeuN). This methodology allowed us to obtain precise estimates of neuronal density at different depths of the cortex, which could be assigned to specific cytoarchitectural fields, thereby significantly extending the scope of previous research. Based on these data, and accurate computational measurements of thickness of the cortex corresponding to different areas, we were also able to estimate the mean numbers of neurons in a column in different parts of the cortex, and the total numbers of neurons in different areas of a marmoset brain hemisphere. To enable future reanalysis, we share the entire data set through a freely available repository (http://www.marmosetbrain.org/cell_density).

### Neuronal density

Based on an analysis of the comprehensive data obtained by Collins et al. (2010) using the isotropic fractionator method, Cahalane et al. (2012) proposed that variations in neuronal density in the primate cortex approximately follow an anterolateral to posteromedial gradient among isocortical areas in the dorsal and lateral surfaces of the primate cortex. However, as acknowledged by these authors, the existence of an overall gradient does not preclude the existence of finer-grained transitions at the borders of areas (which cannot be determined when the isotropic fractionator method is used). For example, in regions consisting of small subdivisions, such as the auditory cortex, this method does not carry enough resolution to separate areas based on differential density. Subsequent work in primates and other species, using stereological methods and a grid strategy (e.g. Charvet et al. 2015, 2016, 2017) has confirmed the existence of a neuronal density gradient, but until the present study there have been no available estimates of neuronal density for the majority of the cytoarchitectural areas in the primate brain.

Our observations support the view that variations in neuronal density in different areas of the marmoset cortex are significant, and follow trends that are consistent with those described in other species. For example, V1 stands out on the basis of its extraordinarily high neuronal density (Rockel et al. 1980), and there is a broad trend for neuronal density to decrease towards the frontal lobe, as previously shown in marsupials (Charvet et al. 2017), rodents (Charvet et al. 2015; Herculano-Houzel et al. 2013), cats (Beaullieu and Collonier 1989; Charvet et al. 2016), sirenians (Charvet et al. 2016), other non-human primates (Cahalane et al. 2012; Charvet et al. 2015; Collins et al. 2010, 2016; Hilgetag et al. 2016) and humans (Ribeiro et al. 2013). We also found that neuronal density in motor and premotor areas is lower than that in either the somatosensory areas or prefrontal areas, in agreement with studies of other non-human primates (Collins et al. 2010, 2016; Hilgetag et al. 2016).

By adopting an approach based on the systematic identification of cytoarchitectural areas, we demonstrate that variations in neuronal density form a complex pattern, with multiple local gradients being apparent (Fig. 12). In visual and somatosensory cortex, the local trends in neuronal density resemble functional hierarchies proposed on the basis of connection patterns and functional properties (Hilgetag et al. 2000; Chaudhuri et al. 2015): neuronal density decreases from posterior to anterior areas (i.e., from V1 and V2 to rostral inferior temporal areas), whereas in somatosensory cortex it decreases from anterior to posterior (from area 3b to PE). These trends are symmetrical to those observed in studies of pyramidal neuron morphology, which reported that the size of neurons (and the number of spines in their dendritic arbor) increases towards anterior visual areas (Elston and Rosa 1997; 1998a, b), but towards posterior somatosensory areas (Elston and Rockland 2002).

**Figure 12.**
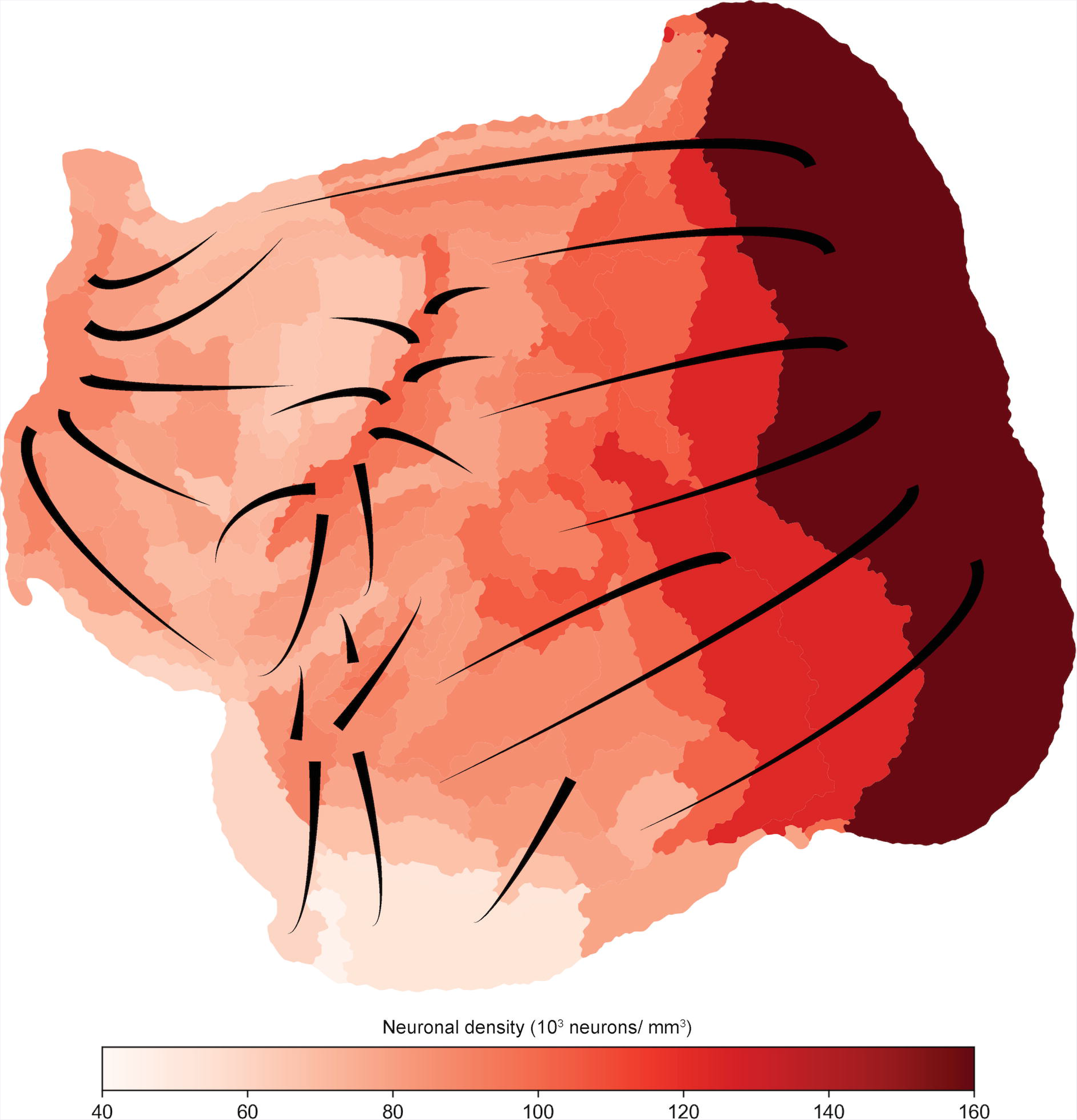
“Unfolded” representation of the marmoset cortex, showing the gradients of neuronal density observed in different regions. The decreasing thickness of the lines indicates reductions of neuronal density along a particular axis.

This finer level of analysis provides some support for the idea that neuronal density decreases in parallel with the functional requirements of different areas, in particular with respect to the need to integrate progressively greater numbers of synaptic inputs, which in turn require greater allocation of space to neuropil (Elston et al. 1999). However, it also raises significant questions in light of this model. For example, auditory areas are currently seen as being organized in a functional hierarchy, from core, to belt, to parabelt (Kaas and Hackett 2000). Our analysis reveals that, in fact, the dominant trend is for the neuronal density to increase along the posterior to anterior axis, in each of these hierarchical levels. Moreover, in the prefrontal cortex, rostral areas (e.g. the frontopolar cortex) are usually seen as being involved in more abstract cognitive functions, in comparison to caudal areas such as the frontal eye fields. Neurons in the former areas are therefore regarded as being the site of wide integration of inputs (Elston 2000), which enable functions related to high-order cognition (Mansouri et al. 2017). However, neuronal density was found to increase towards the frontal pole. These discrepancies suggest that other factors interact with the functional demand for integration of inputs in determining the precise values of neuronal density.

A clue to one of these factors may lie in our finding that area MT (V5), which corresponds to a relatively low level of the dorsal stream hierarchy, is an outlier in terms of having a significantly lower neuronal density in comparison with adjacent, functionally related areas. Here, the functional demand for fast conduction of visual information, which demands extraordinarily high levels of thick, myelinated axons within the cortex, may impose a wider separation between neurons, resulting in lower neuronal density. This factor may also be at the root of the observed gradients among auditory areas (where myelination is higher in caudal areas such as A1 and CM; de la Mothe et al. 2006) and prefrontal areas (where myelination progressively decreases from areas 8 and 45 to area 10; Burman et al. 2006; Burman and Rosa 2009). In addition, factors related to the developmental mechanics in regions of high and low curvature of the cortex, such as the relative expansion and compression of the supragranular layers in regions of high and low convexity (Hilgetag and Barbas 2006), can influence local values of neuronal density. Although we have attempted to select regions of low curvature for measurement, this proved impossible for some of the areas; for example, the external segment of S2 runs along the lip of the lateral sulcus, whereas area PGa/IPa precisely follows the fundus of the superior temporal sulcus.

Finally, part of the observed variation in neuronal density is also likely to result from the application of different genetically influenced developmental programs, which reflect the progressive degree of differentiation of cortical structure in evolution (Sanides 1972). Neuronal density increases gradually as one considers areas progressively distant from the piriform and hippocampal formation; values tend to be low in periallocortical and proisocortical areas of the ventral cingulate, anterior insular and parahippocampal regions, where the cortex is also thinner and less laminated, reflecting the evolutionary gradient proposed by Sanides (1972). However, our data show that, even among these regions, neuronal density is not uniform. For example, posterior limbic areas such as prostriata, area 30 and area 23a have higher neuronal densities than do anterior cingulate and medial prefrontal areas. Likewise, parahippocampal area TH has a higher neuronal density than does the more rostrally located pararhinal cortex (area 36). These observations suggest that an anteroposterior gradient of neuronal density occurs even in limbic areas, in which neurogenesis is completed earlier than in lateral and dorsal isocortical areas (Granger et al. 1995). In summary, our data indicate that neuronal density in the adult cortex represents a compromise between multiple factors, including different genetic programs and activity-dependent processes that reflect the distinct functional demands imposed on different specific neuronal populations in postnatal life. This highlights the need for future studies in which analyses similar to the ones conducted here are applied to animals ranging in age from neonatal to full maturity, which hopefully will help to clarify which differences emerge from the gradual process of circuit maturation in postnatal life (e.g. Bourne and Rosa 2006; Burman et al. 2007).

Our results revealed that the neuronal density in marmoset V1 is approximately more than 3 times higher than that in the amygdalopiriform transition area, which together represent the extremes in our data set. The findings of Hilgetag et al. (2016) suggest that an even steeper gradient may occur in the macaque. For example, in the marmoset the neuronal density in V1 is 3.2 times higher than that in area 35 (entorhinal cortex), whereas in the macaque it is 6.8 times higher. This observation is in line with previous reports suggesting that larger brains tend to show steeper anteroposterior gradients of neuronal density (Schenker et al. 2008; Collins et al. 2010; Semendeferi et al. 2011; Cahalane et al. 2012; Herculano-Houzel et al. 2013; Ribeiro et al. 2013; Charvet et al. 2015; Finlay and Uchiyama 2015).

### Cortical thickness, and number of neurons per column

Using a computational method based on a 3-dimensional reconstruction, we measured cortical thickness in the same brain from which we obtained comprehensive estimates of neuronal density. The distribution of cortical thickness in this individual appears to be, in many aspects, qualitatively similar to that reported for adult macaques (Koo et al. 2012; Calabrese et al. 2015). In both species, regions characterized by thick cortex included parts of the precentral gyrus and operculum, temporal pole, ventrolateral prefrontal cortex, and the ventral precuneus, whereas the tentorial and ventral temporal surfaces (including parahippocampal areas) was formed by relatively thin cortex. In marmosets the intraparietal cortex and dorsal superior temporal regions were also found to form a cluster of areas characterized by thick cortex; however, these areas are located within deep sulci in the macaque brain, preventing comparison with the illustrations presented by Koo et al. (2012) and Calabrese et al. (2015).

Based on data reported by Collins et al. (2010), Calahane et al. (2012), estimated the thickness of the isocortex in the dorsal and lateral surfaces of the brain of other primate species, and reported a statistically significant trend for increasing thickness from posterior to anterior isocortical areas. Our results show that this trend also exists in the marmoset isocortex (Fig. 10), but is relatively subtle (r^2^=0.18), even along a principal axis determined following fitting of a surface (Supplementary Table 2). For example, specific regions, such as the posterior parietal cortex and some of the “dorsal stream” extrastriate areas, stand out as genuine islands of increased thickness among adjacent areas in the caudal half of the brain. Furthermore, thickness is strongly influenced by the degree of local curvature (convexity or concavity) of the cortex, leading to additional significant local variability.

By combining the data on mean neuronal density and cortical thickness, we were also able to estimate the number of neurons within a column of equal sized (1 mm^2^) cross-section, in different cortical areas. This analysis revealed several examples of trends which differ from those suggested by analysis of neuronal density. For example, the core fields (A1, R and RT) proved to have higher number of neurons per unit column in comparison with other auditory areas. Unlike in the analysis of neuronal density, this shows an indication of a functional-hierarchical trend that is shared with visual and somatosensory cortex (where primary areas also show the highest number of neurons per column). Thus, neuronal density and cortical thickness interact in defining the number of neurons involved in local computations, performed by columns of cells in different areas of the adult brain. Among the 3 measures explored (neuronal density, thickness, and number of neurons per column), neuronal density was the one found to vary most consistently across the cortex: using a dataset that excludes limbic areas (Supplementary Table S1), similar to that used in the analyses by Calahane et al. (2012) and Charvet et al. (2015), the coefficient of correlation along the principal axis of variation (r^2^=0.78; Supplementary Figure S5) was higher than those describing the relationships of thickness (r^2^=0.16; Fig. 10) or number of neurons per unit of cortical surface (r^2^= 0.28; Fig. 11).

Charvet et al. (2015) reported that changes in neuron numbers per unit of cortical surface are more pronounced in primates (owl monkey, tamarin and capuchin) than in rodents: whereas in primates ratio between the numbers of neurons in the rostral and caudal poles varied between 1.64 and 2.13, in rodents the corresponding ratio varied between 1.15–1.54. However, this analysis, which was motivated by a neurodevelopmental model based on the gradient in duration of neurogenesis (Rakic 2002), only used areas in the lateral and dorsal surfaces of the brain. In marmosets, across the entire cortex, we found that the number of neurons per 1 mm^2^ unit of cortical surface varied by a factor of 4 between minima in the entorhinal and piriform complexes, and a peak in V1. In humans, the number of neurons per cortical column is reported to vary by a factor of 5 across the anteroposterior axis of the entire cortex (Ribeiro et al., 2013).

### Methodological considerations and possible limitations of the present study

To date, the most comprehensive reports of variations in neuronal density in the primate cortex have been based on the application of the isotropic fractionator method (Herculano-Houzel and Lent 2005) to flat-mounted preparations of the cortex (e.g. Collins et al. 2010, 2016; Turner et al. 2016). Several aspects of the present data are also apparent in results obtained using this method, including the overall anteroposterior gradient of neuronal density, the low neuronal density in motor cortex, and the relatively high density in areas 3b and V1 in comparison with adjacent areas. However, the isotropic fractionator method only allows the precise attribution of neuronal density values to a few areas, namely those which can be approximately delineated by differences in myelination that are visible in non-sectioned flat-mounted preparations. Moreover, it incurs in a loss of information about changes in density across the thickness of the cortex, and potential imprecisions exist due to different (and non-measurable) degrees of compression and expansion of the cortical surface and the introduction of discontinuities during the flat-mounting procedure, as well as the likely inclusion of white matter neurons (which are ubiquitous, but not uniformly distributed in the regions immediately subjacent to layer 6). Another limitation of the isotropic fractionator technique is likely undercounting as a result of tissue dissociation (Charvet et al. 2015).

Although the present method circumvents these problems, it is also important to acknowledge possible factors that limit its precision. Chief among these is the fact that cell counting is manual, repetitive task, which leaves some room for human error and subjectivity. For example, different observers examining Figure 1 may arrive to slightly different numbers, given different interpretations of cell bodies that overlap in the image, whether or not a nucleus is visible, or simple errors such as “missing” a cell. We have attempted to mitigate this by having a quality control procedure that involved two observers working independently, and manual checks of any instances when the cell density profiles showed significant differences. In the vast majority of cases, the differences proved to be genuine, and re-counts resulted in only slight differences (1-2 cells difference within a 150 × 150 μm frame). Thus, imprecision due to human factors is unlikely to have affected the present conclusions. Other possible sources of imprecision could be related to specific features of our workflow. For example, by using high-resolution scanned images, instead of direct observation under the microscope, we were unable to focus up and down, so it is possible that individual cell bodies may have been completely obscured by others. In addition, our decision to deliberately avoid columns where large blood vessels were visible may have slightly inflated the total cell counts. However, adopting an approach based on random allocation of counting strips would carry a significant risk of localized under-counting due to large vessels, which could only be circumvented by considering a much large number of samples. Moreover, it would be difficult to control for effects of differential expansion and shrinkage of vessel and non-vessel tissue during histological processing, which are difficult to estimate. At this point, it is worth emphasizing that all of the potential issues identified above applied equally to the entire sample, which was analyzed using a consistent workflow. Thus, conclusions referring to relative differences between areas would not be significantly affected. The present estimates of V1 neuronal density are in good agreement with those obtained in the most complete stereological study of the macaque cortex to date (Hilgetag et al. 2016), which also used 150 × 150 μm counting frames, and sections of similar thickness (50 vs. 40 μm).

Another type of limitation comes from the fact that cortical areas have cytoarchitectural borders that vary in their degree of definition (Rosa and Tweedale 2005). Thus, whereas estimates obtained for areas with sharp boundaries (e.g. the primary sensory areas, or extrastriate area MT) can be confidently assigned to one functional subdivision of the cortex, for other areas there is a larger margin of error. For example, we obtained lower estimates of neuronal density in the rostral third of inferior temporal area TE3, in comparison with the caudal two-thirds (Supplementary Figure S7). This may reflect imprecision in defining the border of this area, or (in our view, more likely), the fact that a single cytoarchitectural area may contain functional subdivisions that are not evident with the histological methods used. Conversely, some functional areas show regional differences in histology, including the primary motor area (area 4; Burman et al. 2008; Supplementary Figure S7). There is no simple solution to this problem, particularly given that knowledge about the boundaries of cortical continues to evolve. Thus, no single scheme of subdivision, including the one used in the present study (Paxinos et al. 2012) can be regarded as definitive. Acknowledging the fact that some of the present conclusions may need to be refined in light of future developments in this field, we provide the entirety of our sample via an online repository (www.marmosetbrain.org), which will hopefully enable reanalysis. In the meantime, we tried to minimize the effect of local variations and errors of assignment to areas by choosing counting strips which were spread across the extent of each area, while avoiding putative border regions.

The gradual nature of the transition between cortex and white matter, related to the presence of ubiquitous white matter neurons, represents another potential source of imprecision. Our approach was based on visual determination of the lower boundary of the cortex by an experienced neuroanatomist (as illustrated in Fig. 5). However, this determination is fundamentally subjective, and this problem is exacerbated in highly convex regions, in which the white matter may contain scattered neurons even hundreds of micrometers below the estimated layer 6 boundary (e.g. areas TEO, A1 and 6M, in Fig. 5). Inclusion of what we considered white matter neurons would have resulted in lower mean neuronal densities, and slightly higher numbers of neurons per column.

Finally, it is important to acknowledge that the fact that only one individual’s brain was extensively sampled. Our less extensive measurements obtained in 2 additional animals provide some reassurance that the features we describe are likely representative of both male and female marmosets, but do not address important questions such as potential age-related changes (which are likely; Amlien et al. 2016), and differences between hemispheres. Future developments in automated image analysis (which may enable neuronal counting with a similar precision currently achieved by humans) and histological processing pipelines (which may reduce the number of histological artifacts likely to affect measurements; Lin et al. 2018) will likely facilitate studies involving larger sample sizes.

### Future directions

It has been argued that neuronal density is a strong predictor of patterns of cortico-cortical connectivity. Hilgetag et al. (2016), building on earlier work on the connections of frontal areas (e.g. Dombrowski et al. 2001; Hilgetag and Grant 2010), reported that cytoarchitectural similarity is a reliable predictor of the existence and strength of connections between any two areas. These authors also found that cortico-cortical laminar patterns of connections correlate with structurally similarity: dissimilar areas tend to be linked by connections that show strong laminar biases (i.e. terminations and origins located in predominantly either supra- or infragranular layers), whereas similar areas tend to show more balanced origins and terminations throughout the depth of the cortex. These conclusions were based on quantitative analysis of mean neuronal density in a subset of cortical areas, combined with qualitative assessments of cytoarchitectural type. A more comprehensive exploration of structural relationships between cytoarchitecture and connectivity, based on statistical analysis, will likely require a fully quantitative approach. Moreover, incorporation of a finer level of knowledge, in the form of estimates throughout the depth of the cortex, may reveal additional principles, which are not transparent to models that use average density as a single dimension. When combined with current efforts to obtain a comprehensive understanding of the cortico-cortical connectivity in the marmoset (Okano and Mitra 2015; Majka et al. 2016; Liu et al. 2018), this will provide unique new opportunities to decipher the logic of cortical connectivity. In addition, future work using the present methods may allow determination of the densities of different neuronal subtypes in the marmoset brain, allowing a greater level of insight on the neuronal computations being performed by different areas (Kim et al. 2017; Wang and Yang 2018).

## Acknowledgements

The authors thank Jonathan Chan, Cecilia Cranfield and Natalia Jermakow for assistance in many aspects of this project, Shi Bai for developing the computational infrastructure that enables us to share the original data sets used in this analysis, and Rowan Tweedale for expert editing of an earlier version of the manuscript. We also thank the Monash Histology Platform for slide scanning, and the Monash e-Research Centre for access to the MASSIVE computer infrastructure, used for 3-D reconstruction and registration of the cortex. Supported by the Australian Research Council (grants DP140101968 and CE140100007), National Health and Medical Research Council (grant 1122220) and the International Neuroinformatics Coordinating Facility (INCF) seed funding scheme.

## References

Angelucci A, Rosa MGP. 2015. Resolving the organization of the third tier visual cortex in primates: a hypothesis-based approach. Vis Neurosci. 32: E010.

Amlien IK, Fjell AM, Tamnes CK, Grydeland H, Krogsrud SK, Chaplin TA, Rosa MGP, Walhovd KB. 2016. Organizing principles of human cortical development--thickness and area from 4 to 30 years: insights from comparative primate neuroanatomy. Cereb Cortex 26: 257–267.

Atapour N, Worthy KH, Lui LL, Yu HH, Rosa MGP. 2017. Neuronal degeneration in the dorsal lateral geniculate nucleus following lesions of primary visual cortex: comparison of young adult and geriatric marmoset monkeys. Brain Struct Funct 222: 3283–3293.

Bakola S, Burman KJ, Rosa MGP. 2015. The cortical motor system of the marmoset monkey (Callithrix jacchus). Neurosci Res 93: 72–81.

Baylis GC, Rolls ET, Leonard CM. 1987. Functional subdivisions of the temporal lobe neocortex. J Neurosci 7: 330–342.

Beaullieu C, Collonier M. 1989. Effects of the richness of the environment on six different cortical areas of the cat cerebral cortex. Brain Res 495: 382–386.

Bourne JA, Rosa MGP. 2006. Hierarchical development of the primate visual cortex, as revealed by neurofilament immunoreactivity: early maturation of the middle temporal area (MT). Cereb Cortex 16: 405–414.

Bunk EC, Stelzer S, Hermann S, Schäfers M, Schlatt S, Schwamborn JC. 2011. Cellular organization of adult neurogenesis in the common marmoset. Aging Cell 10: 28–38.

Burman KJ, Palmer SM, Gamberini M, Rosa MGP. 2006. Cytoarchitectonic subdivisions of the dorsolateral frontal cortex of the marmoset monkey (Callithrix jacchus), and their projections to dorsal visual areas. J Comp Neurol 495: 149–172.

Burman KJ, Lui LL, Rosa MGP, Bourne JA. 2007. Development of non-phosphorylated neurofilament protein expression in neurones of the New World monkey dorsolateral frontal cortex. Eur J Neurosci 25: 1767–1779.

Burman KJ, Palmer SM, Gamberini M, Spitzer MW, Rosa MGP. 2008. Anatomical and physiological definition of the motor cortex of the marmoset monkey. J Comp Neurol 506: 860–876.

Burman KJ, Rosa MGP. 2009. Architectural subdivisions of medial and orbital frontal cortices in the marmoset monkey (CaJJithrixjacchus). J Comp Neurol 514: 11–29.

Burman KJ, Bakola S, Richardson KE, Reser DH, Rosa MGP. 2014a. Patterns of cortical input to the primary motor area in the marmoset monkey. J Comp Neurol 522: 811–843.

Burman KJ, Bakola S, Richardson KE, Reser DH, Rosa MGP. 2014b. Patterns of afferent input to the caudal and rostral areas of the dorsal premotor cortex (6DC and 6DR) in the marmoset monkey. J Comp Neurol 522: 3683–3716.

Burman KJ, Bakola S, Richardson KE, Yu HH, Reser DH, Rosa MGP. 2015. Cortical and thalamic projections to cytoarchitectural areas 6Va and 8C of the marmoset monkey: connectionally distinct subdivisions of the lateral premotor cortex. J Comp Neurol 523: 1222–1247.

Cahalane DJ, Charvet CJ, Finlay BL. 2012. Systematic, balancing gradients in neuron density and number across the primate isocortex. Front Neuroanat 6: 28.

Calabrese E, Badea A, Coe CL, Lubach GR, Shi Y, Styner MA, Johnson GA. 2015. A diffusion tensor MRI atlas of the postmortem rhesus macaque brain. Neuroimage 117: 408–416

Chaplin TA, Yu HH, Rosa MGP. 2013. Representation of the visual field in the primary visual area of the marmoset monkey: magnification factors, point-image size, and proportionality to retinal ganglion cell density. J Comp Neurol 521: 1001–1019.

Charvet CJ, Cahalane DJ, Finlay BL. 2015. Systematic, cross-cortex variation in neuron numbers in rodents and primates. Cereb Cortex 25: 147–160.

Charvet CJ, Reep RL, Finlay BL. 2016. Evolution of cytoarchitectural landscapes in the mammalian isocortex: Sirenians (Trichechus manatus) in comparison with other mammals. J Comp Neurol 524: 772–782.

Charvet CJ, Stimpson CD, Kim YD, Raghanti MA, Lewandowski AH, Hof PR, Gómez-Robles A, Krienen FM, Sherwood CC. 2017. Gradients in cytoarchitectural landscapes of the isocortex: Diprotodont marsupials in comparison to eutherian mammals. J Comp Neurol 525: 1811–1826.

Chaudhuri R, Knoblauch K, Gariel MA, Kennedy H, Wang XJ. 2015. A large-scale circuit mechanism for hierarchical dynamical processing in the primate cortex. Neuron. 2015 Oct 21;88(2):419–31.

Collins CE, Airey DC, Young NA, Leitch DB, Kaas JH. 2010. Neuron densities vary across and within cortical areas in primates. Proc Natl Acad Sci U S A 107: 15927–15932.

Collins CE. 2011. Variability in neuron densities across the cortical sheet in primates. Brain Behav Evol 78: 37–50.

Collins CE, Turner EC, Sawyer EK, Reed JL, Young NA, Flaherty DK, Kaas JH. 2016. Cortical cell and neuron density estimates in one chimpanzee hemisphere. Proc Natl Acad Sci U S A 113: 740–745.

de la Mothe LA, Blumell S, Kajikawa Y, Hackett TA. 2006. Cortical connections of the auditory cortex in marmoset monkeys: core and medial belt regions. J Comp Neurol 496: 27–71.

Dombrowski SM, Hilgetag CC, Barbas H. 2001. Quantitative architecture distinguishes prefrontal cortical systems in the rhesus monkey. Cereb Cortex 11: 975–988.

Dorph-Petersen KA, Caric D, Saghafi R, Zhang W, Sampson AR, Lewis DA. 2009. Volume and neuron number of the lateral geniculate nucleus in schizophrenia and mood disorders. Acta Neuropathol 117: 369–384.

Eliades SJ, Miller CT. 2017. Marmoset vocal communication: Behavior and neurobiology. Dev Neurobiol 77: 286–299.

Elston GN. 2000. Pyramidal cells of the frontal lobe: all the more spinous to think with. J Neurosci 20: RC95.

Elston GN, Rockland KS. 2002. The pyramidal cell of the sensorimotor cortex of the macaque monkey: phenotypic variation. Cereb Cortex 12: 1071–1078.

Elston GN, Rosa MGP. 1997. The occipitoparietal pathway of the macaque monkey: comparison of pyramidal cell morphology in layer III of functionally related cortical visual areas. Cereb Cortex 7: 432–452.

Elston GN, Rosa MGP. 1998a. Morphological variation of layer III pyramidal neurones in the occipitotemporal pathway of the macaque monkey visual cortex. Cereb Cortex 8: 278–294.

Elston GN, Rosa MGP. 1998b. Complex dendritic fields of pyramidal cells in the frontal eye field of the macaque monkey: comparison with parietal areas 7a and LIP. Neuroreport 9: 127–131.

Elston GN, Tweedale R, Rosa MGP. 1999. Cortical integration in the visual system of the macaque monkey: large-scale morphological differences in the pyramidal neurons in the occipital, parietal and temporal lobes. Proc R Soc Lond B Biol Sci 266:1367–1374.

Finlay BL, Uchiyama R. 2015. Developmental mechanisms channeling cortical evolution. Trends Neurosci 38: 69–76.

Gallyas F.1979. Silver staining of myelin by means of physical development. Neurol Res 1:203–209.

Gittins R, Harrison PJ. 2004. Neuronal density, size and shape in the human anterior cingulate cortex: a comparison of Nissl and NeuN staining. Brain Res Bull 63: 155–160.

Granger B, Tekaia F, Le Sourd AM, Rakic P, Bourgeois JP. 1995. Tempo of neurogenesis and synaptogenesis in the primate cingulate mesocortex: comparison with the neocortex. J Comp Neurol 360: 363–376.

Hagan MA, Rosa MGP, Lui LL. 2017. Neural plasticity following lesions of the primate occipital lobe: The marmoset as an animal model for studies of blindsight. Dev Neurobiol 77: 314–327.

Herculano-Houzel S, and Lent R. 2005. Isotropic fractionator: a simple, rapid method for the quantification of total cell and neuron numbers in the brain. J. Neurosci 25: 2518–2521.

Herculano-Houzel S, Watson C, Paxinos G. 2013. Distribution of neurons in functional areas of the mouse cerebral cortex reveals quantitatively different cortical zones. Front Neuroanat 7: 35.

Hilgetag CC, O’Neill MA, Young MP. 2000. Hierarchical organization of macaque and cat cortical sensory systems explored with a novel network processor. Philos Trans R Soc Lond B Biol Sci 355:71–89.

Hilgetag CC, Barbas H. 2006. Role of mechanical factors in the morphology of the primate cerebral cortex. PLoS Comput Biol 2: e22.

Hilgetag CC, Grant S. 2010. Cytoarchitectural differences are a key determinant of laminar projection origins in the visual cortex. Neuroimage 51: 1006–1017.

Hilgetag CC, Medalla M, Beul SF, Barbas H. 2016. The primate connectome in context: Principles of connections of the cortical visual system. Neuroimage 134: 685–702.

Huffman KJ, Krubitzer L. 2001. Area 3a: topographic organization and cortical connections in marmoset monkeys. Cereb Cortex 11: 849–867.

Hustler JJ, Lee DG, Porter KK. 2005. Comparative analysis of cortical layering and supragranular layer enlargement in rodent carnivore and primate species. Brain Res 1052: 71–81.

Jeffs J, Federer F, Ichida JM, Angelucci A. 2013. High-resolution mapping of anatomical connections in marmoset extrastriate cortex reveals a complete representation of the visual field bordering dorsal V2. Cereb Cortex 23: 1126–1147.

Jeffs J, Federer F, Angelucci A. 2015. Corticocortical connection patterns reveal two distinct visual cortical areas bordering dorsal V2 in marmoset monkey. Vis Neurosci 32: E012.

Jones SE, Buchbinder BR, Aharon I. 2000. Three-dimensional mapping of cortical thickness using Laplace’s equation. Hum Brain Mapp. 11: 12–32

Kaas JH, Hackett TA. 2000. Subdivisions of auditory cortex and processing streams in primates. Proc Natl Acad Sci U S A 97:11793–11799.

Kim Y, Yang GR, Pradhan K, Venkataraju KU, Bota M, García Del Molino LC, Fitzgerald G, Ram K, He M, Levine JM, Mitra P, Huang ZJ, Wang XJ, Osten P. 2017. Brain-wide maps reveal stereotyped cell-type-based cortical architecture and subcortical sexual dimorphism. Cell 171: 456–469.

Koo BB, Schettler SP, Murray DE, Lee JM, Killiany RJ, Rosene DL, Kim DS, Ronen I. 2012. Age-related effects on cortical thickness patterns of the rhesus monkey brain. Neurobiol Aging 33(1):200.e23–31.

Krubitzer LA, Kaas JH. 1990. The organization and connections of somatosensory cortex in marmosets. J Neurosci 10: 952–974.

Lin MK, Takahashi YS, Huo B-X, Hanada M, Nagashima J, Hata J, Tolpygo AS, Ram K, Lee BC, Miller MI, Rosa MGP, Sasaki E, Iriki A, Okano H, Mitra PP. 2018. A high-throughput neurohistological pipeline for brain-wide mesoscale connectivity mapping of the common marmoset. bioRxiv.315804; preprint in open access repository.

Liu C, Ye FQ, Yen CC, Newman JD, Glen D, Leopold DA, Silva AC. 2018. A digital 3D atlas of the marmoset brain based on multi-modal MRI. Neuroimage 169: 106–116.

Majka P, Chaplin TA, Yu HH, Tolpygo A, Mitra PP, Wójcik DK, Rosa MGP. 2016. Towards a comprehensive atlas of cortical connections in a primate brain: Mapping tracer injection studies of the common marmoset into a reference digital template. J Comp Neurol 524: 2161–2181.

Mansouri FA, Koechlin E, Rosa MGP, Buckley MJ. 2017. Managing competing goals-a key role for the frontopolar cortex. Nat Rev Neurosci 18: 645–657.

Miller CT, Freiwald WA, Leopold DA, Mitchell JF, Silva AC, Wang X. 2016. Marmosets: A neuroscientific model of human social behavior. Neuron 90: 219–233.

Mitchell JF, Leopold DA. 2015. The marmoset monkey as a model for visual neuroscience. Neurosci Res 93: 20–46.

Mortazavi F, Wang X, Rosene DL, Rockland KS. 2016. White matter neurons in young adult and aged rhesus monkey. Front Neuroanat 10: 15.

Mortazavi F, Romano SE, Rosene DL, Rockland KS. 2017. A survey of white matter neurons at the gyral crowns and sulcal depths in the rhesus monkey. Front Neuroanat 11: 69.

Mullen RJ, Buck CR, Smith AM. 1992. NeuN, a neuronal specific nuclear protein in vertebrates. Development 116: 201–211.

Oikonomidis L, Santangelo AM, Shiba Y, Clarke FH, Robbins TW, Roberts AC. 2017. A dimensional approach to modeling symptoms of neuropsychiatric disorders in the marmoset monkey. Dev Neurobiol 77: 328–353.

Okano H, Kishi N. 2017. Investigation of brain science and neurological/psychiatric disorders using genetically modified non-human primates. Curr Opin Neurobiol 50: 1–6.

Okano H, Mitra P. 2015. Brain-mapping projects using the common marmoset. Neurosci Res 93: 3–7.

Padberg J, Franca JG, Cooke DF, Soares JG, Rosa MGP, Fiorani M Jr, Gattass R, Krubitzer L. 2007. Parallel evolution of cortical areas involved in skilled hand use. J Neurosci 27: 10106–10115.

Palmer SM, Rosa MGP. 2006a. A distinct anatomical network of cortical areas for analysis of motion in far peripheral vision. Eur J Neurosci 24: 2389–2405.

Palmer SM, Rosa MGP. 2006b. Quantitative analysis of the corticocortical projections to the middle temporal area in the marmoset monkey: evolutionary and functional implications. Cereb Cortex 16: 1361–1375.

Passarelli L, Rosa MGP, Bakola S, Gamberini M, Worthy KH, Fattori P, Galletti C. 2018. Uniformity and diversity of cortical projections to precuneate areas in the macaque monkey: what defines area PGm? Cereb Cortex 28: 1700–1717.

Paxinos G, Watson C, Petrides M, Rosa MGP, Tokuno H. 2012. The Marmoset Brain in Stereotaxic Coordinates. New York: Academic Press.

Rakic, P. 2002. Pre-and post-developmental neurogenesis in primates. Clin Neurosci Res 2, 29–39.

Reser DH, Burman KJ, Richardson KE, Spitzer MW, Rosa MGP. 2009. Connections of the marmoset rostrotemporal auditory area: express pathways for analysis of affective content in hearing. Eur J Neurosci 30: 578–592.

Reser DH, Burman KJ, Yu HH, Chaplin TA, Richardson KE, Worthy KH, Rosa MGP. 2013. Contrasting patterns of cortical input to architectural subdivisions of the area 8 complex: a retrograde tracing study in marmoset monkeys. Cereb Cortex 23: 1901–1922.

Ribeiro PF, Ventura-Antunes L, Gabi M, Mota B, Grinberg LT, Farfel JM, Ferretti-Rebustini RE, Leite RE, Filho WJ, Herculano-Houzel S. 2013. The human cerebral cortex is neither one nor many: neuronal distribution reveals two quantitatively different zones in the gray matter, three in the white matter, and explains local variations in cortical folding. Front Neuroanat 7: 28.

Rockel AJ, Hiorns RW, Powell TPS. 1980. The basic uniformity in structure of the neocortex. Brain 103:221–244.

Rosa MGP, Elston GN. 1998. Visuotopic organisation and neuronal response selectivity for direction of motion in visual areas of the caudal temporal lobe of the marmoset monkey (Callithrix jacchus): middle temporal area, middle temporal crescent, and surrounding cortex. J Comp Neurol 393: 505–527.

Rosa MGP, Fritsches KA, Elston GN. 1997. The second visual area in the marmoset monkey: visuotopic organisation, magnification factors, architectonical boundaries, and modularity. J Comp Neurol 387: 547–567.

Rosa MGP, Palmer SM, Gamberini M, Tweedale R, Piñon MC, Bourne JA. 2005. Resolving the organization of the New World monkey third visual complex: the dorsal extrastriate cortex of the marmoset (Callithrix jacchus). J Comp Neurol 483: 164–191.

Rosa MGP, Palmer SM, Gamberini M, Burman KJ, Yu HH, Reser DH, Bourne JA, Tweedale R, Galletti C. 2009. Connections of the dorsomedial visual area: pathways for early integration of dorsal and ventral streams in extrastriate cortex. J Neurosci 29: 4548–4563.

Rosa MGP, Schmid LM. 1995. Visual areas in the dorsal and medial extrastriate cortices of the marmoset. J Comp Neurol. 359: 272–299.

Rosa MGP, Soares JGM, Chaplin TA, Majka P, Bakola S, Phillips KA, Reser DH, Gattass R. 2018. Cortical afferents of area 10 in Cebus monkeys: implications for the evolution of the frontal pole. Cereb Cortex, in press. DOI: 10.1093/cercor/bhy044

Rosa MGP, Tweedale R. 2000. Visual areas in lateral and ventral extrastriate cortices of the marmoset monkey. J Comp Neurol 422: 621–651.

Rosa MGP, Tweedale R. 2005. Brain maps, great and small: lessons from comparative studies of primate visual cortical organization. Philos Trans R Soc Lond B Biol Sci 360: 665–691.

Saleem KS, Kondo H, Price JL. 2008. Complementary circuits connecting the orbital and medial prefrontal networks with the temporal, insular, and opercular cortex in the macaque monkey. J Comp Neurol 506: 659–693.

Sanides, F. 1972. Representation in the cerebral cortex and its areal lamination patterns. In: The Structure and Function of Nervous Tissue, ed. Bourne, G. H. Academic Press.

Schenker NM, Buxhoeveden DP, Blackmon WL, Amunts K, Zilles K, Semendeferi K. 2008. A comparative quantitative analysis of cytoarchitecture and minicolumnar organization in Broca’s area in humans and great apes. J Comp Neurol 510: 117–128.

Schmitz C, Hof PR. 2005. Design-based stereology in neuroscience. Neuroscience 130: 813–831.

Schroeder W. 2006. Visualization Toolkit: An Object-Oriented Approach to 3D Graphics, 4th Edition. Clifton Park, NY: Kitware Inc.

Semendeferi K, Teffer K, Buxhoeveden DP, Park MS, Bludau S, Amunts K, Travis K, Buckwalter J. 2011. Spatial organization of neurons in the frontal pole sets humans apart from great apes. Cereb Cortex 21: 1485–1497.

Sherwood CC, Raghanti MA, Stimpson CD, Bonar CJ, de Sousa AA, Preuss TM, Hof PR. 2007. Scaling of inhibitory interneurons in areas V1 and V2 of anthropoid primates as revealed by calcium-binding protein immunohistochemistry. Brain Behav Evol 69: 176–195.

Silva AC. 2017. Anatomical and functional neuroimaging in awake, behaving marmosets. Dev Neurobiol. 77: 373–389.

Silverman MS, Tootell RB. 1987. Modified technique for cytochrome oxidase histochemistry: increased staining intensity and compatibility with 2-deoxyglucose autoradiography. J Neurosci Methods 19: 1–10.

Solomon SG, Rosa MGP. 2014. A simpler primate brain: the visual system of the marmoset monkey. Front. Neural Circuits 8: 96.

Turner EC, Young NA, Reed JL, Collins CE, Flaherty DK, Gabi M, Kaas JH. 2016. Distributions of cells and neurons across the cortical sheet in Old World macaques. Brain Behav Evol 88: 1–13.

Waehnert MD, Dinse J, Weiss M, Streicher MN, Waehnert P, Geyer S, Turner R, Bazin PL. 2013. Anatomically motivated modeling of cortical laminae. Neuroimage 93: 210–220.

Wang XJ, Yang GR. 2018. A disinhibitory circuit motif and flexible information routing in the brain. Curr Opin Neurobiol 49: 75–83.

Woodward A, Hashikawa T, Maeda M, Kaneko T, Hikishima K, Iriki A, Okano H, Yamaguchi Y. 2018. The Brain/MINDS 3D digital marmoset brain atlas. Sci Data 5: 180009.

Young NA, Collins CE, Kaas JH. 2013. Cell and neuron densities in the primary motor cortex of primates. Front Neural Circuits 7: 30.

Yushkevich PA, Piven J, Hazlett HC, Smith RG, Ho S, Gee JC, Gerig G. 2006. User-guided 3D active contour segmentation of anatomical structures: significantly improved efficiency and reliability. NeuroImage 31: 1116–1128.

